# Rugged fitness landscapes minimize promiscuity in the evolution of transcriptional repressors

**DOI:** 10.1101/2022.10.25.513693

**Authors:** Anthony T. Meger, Matthew A. Spence, Mahakaran Sandhu, Colin J. Jackson, Srivatsan Raman

## Abstract

How a protein’s function influences the shape of its fitness landscape, smooth or rugged, is a fundamental question in evolutionary biochemistry. Smooth landscapes arise when incremental mutational steps lead to a progressive change in function, as commonly seen in enzymes and binding proteins. On the other hand, rugged landscapes are poorly understood because of the inherent unpredictability of how sequence changes affect function. Here, we experimentally characterize the entire sequence phylogeny, comprising 1158 extant and ancestral sequences, of the DNA-binding domain (DBD) of the LacI/GalR transcriptional repressor family. Our analysis revealed an extremely rugged landscape with rapid switching of specificity even between adjacent nodes. Further, the ruggedness arises due to the necessity of the repressor to simultaneously evolve specificity for asymmetric operators and disfavors potentially adverse regulatory crosstalk. Our study provides fundamental insight into evolutionary, molecular, and biophysical rules of genetic regulation through the lens of fitness landscapes.

## INTRODUCTION

A central question in molecular evolution is how proteins acquire novel functions, such as new binding specificities, via mutation and selection. The sequence-fitness landscape is a valuable construct that has allowed us to understand and visualize the complex relationship between evolutionary sequence changes and protein function.^1–5^ The topology of a fitness landscape largely reflects the nature of the evolutionary process: “smooth” landscapes, with small numbers of discrete but connected peaks across which new activities can gradually evolve via additive and predictable mutational steps, are relatively well understood, with many enzymes and binding proteins as examples.^2,6–10^ The molecular basis for the smoothness of these transitions can be partly explained by the modulation of the conformational dynamics of proteins, in which mutations can gradually shift the conformational equilibrium towards conformations better suited to the new activity.^11–13^ In contrast, our understanding and characterization of “rugged” fitness landscapes, in which mutations tend to have unpredictable epistatic effects that result in fitness landscapes consisting of multiple peaks separated by many non-functional sequences (“valleys”), is less well developed.^14–16^ One intriguing hypothesis is that the sequence-fitness landscapes of certain proteins are inherently defined by the function and fold.^17^ This partially deterministic view has important implications for evolution of new activities (such as drug^17,18^ or vaccine resistance^19^) and protein engineering. ^20,21^

To understand and characterize the role of rugged fitness landscapes in molecular evolution, we need to comprehensively map sequence-function relationships over large and diverse sequence spaces that span the full evolutionary history of protein families. While directed evolution and deep mutational scanning (DMS) experiments have provided profound insight into molecular evolution at the level of single mutational steps, they seldom explore the large evolutionary timescales and the full diversity of sequence space within protein families.^4,7^ In contrast, ancestral sequence reconstruction (ASR) can sample large spans of sequence space through the computational reconstruction of ancestral species from a phylogenetic tree and a sequence evolution model.^22,23^ However, ASR studies generally characterize a relatively small number of divergent ancestral sequences, which makes it difficult to deconvolute adaptive and neutral mutations and understand the stepwise additive and context-specific (epistatic) effects of mutations.^24^

In contrast to enzymes, which have been shown to often evolve across relatively smooth fitness landscapes in which promiscuous activities can be gradually optimized to become the primary function,^11–13^ transcription factors likely display a more rugged fitness landscape as they are generally defined by high DNA specificity, and incremental mutations leading to promiscuity could be evolutionarily disfavored. The lac repressor (LacI), which belongs to the LacI/GalR family (LGF) of prokaryotic gene regulators,^25^ is a well-studied model for DNA recognition and allostery.^26^ Proteins of the LGF show remarkable diversity in their amino acid sequences and their DNA recognition. While much is known about the operator specificity of *Escherichia coli* LacI (EcLacI),^27–31^ our understanding of the sequence-fitness landscape of DNA-binding specificity in the wider LGF, the historical evolutionary trajectory of operator recognition and the molecular mechanisms that underpin the evolution of DNA-specificity in proteins remains incomplete. The LGF is thus a compelling system to study ruggedness in sequence-fitness landscapes and how the combination of structure and function dictate the evolutionary dynamics and selective pressures exerted on these regulators.

In this study, we experimentally characterize a complete phylogenetic tree (1158 extant and ancestral sequences) of the LGF and perform DMS on extant EcLacI to reveal the sequence-fitness landscape of DNA specificity for the *E. coli* Lac operator. We find the landscape to be extremely rugged due to high levels of epistasis, with most sequences having no affinity for the *E. coli* lac operator sequence. However, the screen unearthed dozens of functional repressors from distinct phylogenetic clades, with as many as 32/60 amino acid substitutions in the DNA binding domain (DBD) compared to the EcLacI DBD. Analysis of the local evolutionary landscape within clades shows gain/loss of function between adjacent nodes, indicating rapid switches of specificity, which may be beneficial for developing orthogonal genetic regulation. The ruggedness of this fitness landscape may therefore be a characteristic feature of regulators and an essential functional requirement of high specificity and orthogonality. The molecular basis for this observation is revealed through *in vitro* binding assays and simulations, which show that ruggedness of the LGF fitness landscape arises in part due to the necessity for regulators to simultaneously evolve specificity for either DNA half-site in asymmetric operators and explains why this fold and operator structure has been evolutionarily selected for genetic regulation. This study provides new fundamental insight into nature’s design of genetic regulation, evolutionary rules of protein-DNA recognition, the biophysical mechanism of transcription factor allostery, and how structure and function combine to determine the sequence-fitness landscapes of proteins.

## RESULTS

### Synthesis and characterization of a complete phylogeny of LGF DNA-binding domains

The *E. coli* lac operator sequence has emerged over a ~3-billion-year evolutionary history. To generate a set of DBDs that spans the full evolutionarily accessible sequence space of the LGF, we performed phylogenetic inference to reconstruct 577 ancestral sequences from a dataset of 581 extant EcLacI homologs (**Fig. 1a**). Consistent with previous phylogenetic studies,^32^ our analysis reconstructs the family as three major lineages. When rooted at the phylogenetic midpoint, these include the LacI-clade as the most ancestral, a single common ancestor giving rise to descendent clades comprising the catabolite control protein (CcpA/RegA), ribose, maltose, sucrose, galactose, arbutin/salicin, purine and cytosine regulators (RbsR, MalR, SacR/ScrR, GalR/GalS, AscG, PurR and CytR, respectively), and an uncharacterized lineage of LacI-like proteins (**Fig. 1a**). On average, the mean posterior probability of each ancestral DBD was 93%, indicating strong statistical support across the full dataset (**Supplementary Fig. 1a**). Likewise, phylogenetic branch supports were consistently high across the topology (**Supplementary Fig. 1b**), and the approximately unbiased test failed to reject the topology we present in Fig. 1 among 9 other tree-search replicates (P-value=0.682)^33^ (**Supplementary Fig. 2**).

**Fig. 1.**
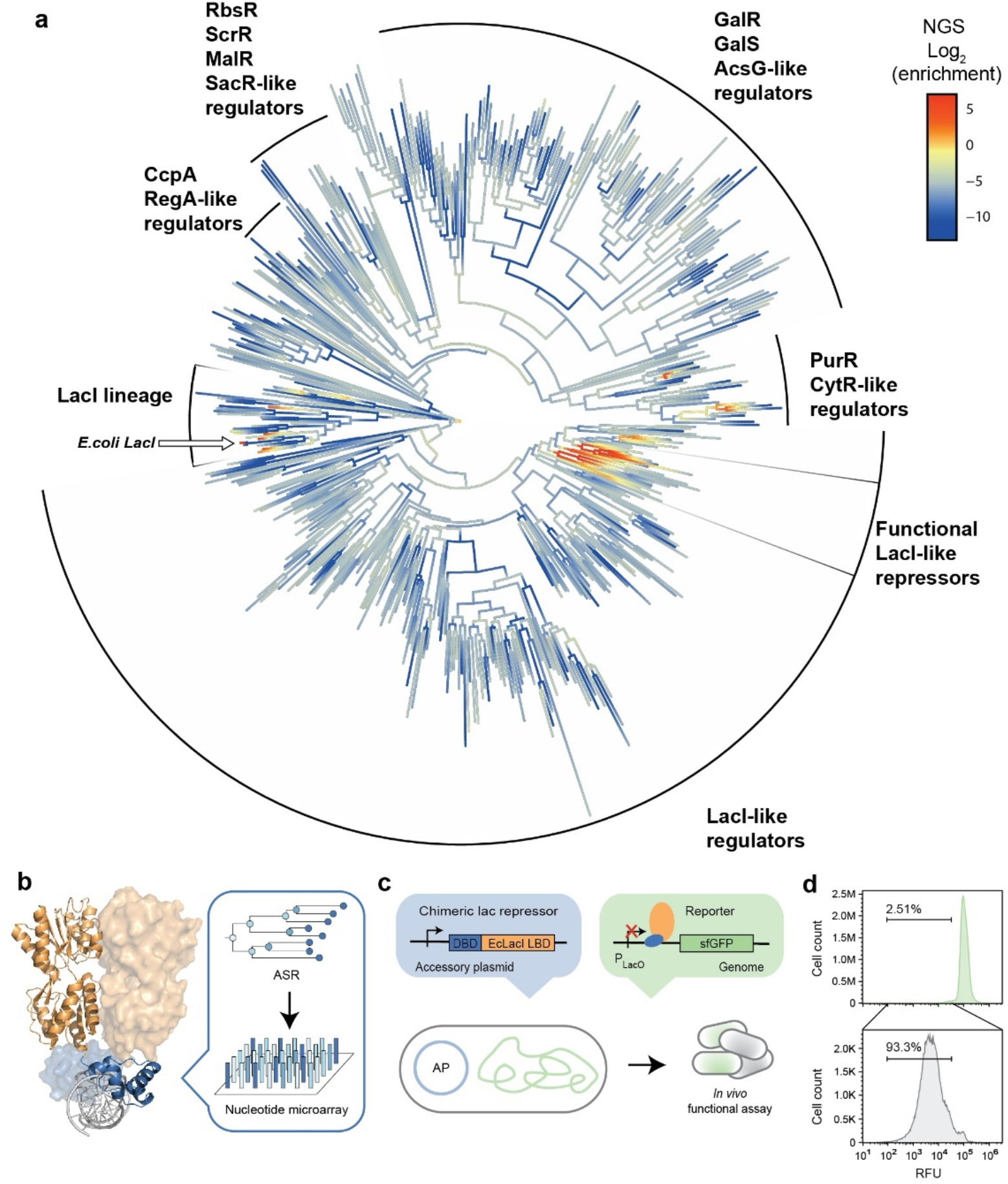
Evolution of LacO recognition. (**a**), Phylogenetic inference of the LacI/GalR family of prokaryotic transcription factors. 577 sequences were ancestrally reconstructed using from a dataset of 581 extant LacI homologs (**Supplementary Fig. 1**, see “methods”). Color gradient shows average repression scores from replicate (*n*=2) high-throughput screens (**Supplementary Fig. 4**). (**b**), Structure (PDB ID: 1EFA) of dimeric extant *E. coli* lac repressor bound to LacO (left). LBD (orange), DBD (blue), and LacO (gray) are shown in cartoon representation. Each extant and ancestral DBD was generated using multiplex micro-array DNA synthesis (right). (**c**), *In vivo* characterization of repressor function. DBDs were fused to the *E. coli* LacI LBD and constitutively expressed on an accessory plasmid. A reporter construct consisting of the gene encoding sfGFP under transcriptional control of LacO was integrated into the genome of E. coli to assay repressor function. (**d**), Flow cytometry of the pre-selected phylogenetic DBD library (top) and after two sequential sorts to isolate repression competent DBDs (bottom). The indicated gate was used for selection by FACS (**Supplementary Fig. 4**). RFU, relative fluorescence units.

One limitation of ASR is that, while it is possible to generate hundreds of ancestral sequences computationally, only a handful (<10) of evolutionarily distant nodes are typically characterized experimentally. Thus, the incremental functional adaptations associated with the full evolutionary trajectory can remain obscured by neutral variation. Here, we synthesized the DBDs of all 577 reconstructed ancestors, as well as all 581 extant sequences used in our phylogeny, generating a dataset of 1158 diverse DBDs that fully covers the phylogenetic tree constructed in this work (**Fig. 1a**). We used chip-based oligonucleotides to synthesize the DBD sequences and cloned them into a plasmid library to encode chimeric variants consisting of DBDs fused to the ligand-binding domain (LBD) of EcLacI, which is induced by allolactose/IPTG (**Fig. 1b**). We constructed chimeras with an invariant LBD for several reasons: (i) by studying the ancestral DBD in the context of the EcLacI LBD, we can be confident that the regulatory mode of the chimeras will match that of the extant EcLacI and will repress in the absence of an allosteric effector;^34^ (ii) since the inducer of the EcLacI LBD is known, we can study the allosteric communication between LBD and DBD in the chimeras; (iii) we can ensure that the experimental conditions we used were conducive to repression, giving accurate insight on exclusively DNA specificity in the DBD. Deep sequencing the plasmid library confirmed 100% coverage of the DBD library with minimal skew i.e., all 1158 DBDs were present (**Supplementary Fig. 3**).

To characterize the affinity of the DBDs within the phylogeny to the *E. coli* lac operator and their allosteric activation by IPTG, we used a cell-based pooled screening strategy (**Fig. 1c**). We inserted a single-copy GFP cassette under the control of the lac operator (LacO) into the *E. coli* genome. Variants that can bind to LacO with sufficient affinity repress GFP expression, while those that cannot produce high GFP expression. Using fluorescence-activated cell sorting (FACS), repression-competent variants are enriched by sorting low GFP cells. Only a small fraction (2.5-2.6%) of low-fluorescence cells were observed via flow cytometry, indicating that relatively few extant and ancestral DBDs in this library repress LacO (**Fig. 1d**). The subset of repression competent variants was enriched to 92.7-93.3% of the total population after two rounds of sorting (**Supplementary Fig. 4**). The pre- and post-sorted libraries were sequenced to estimate the enrichment ratios. A higher enrichment ratio implies greater GFP repression and is a proxy for protein-DNA affinity. The enrichment ratios were highly correlated (R^2^=0.99) between independent replicates (**Supplementary Fig. 5**). We found 15 ancestral and 7 extant DBDs that can repress GFP from the 1158-member library. These 22 enriched DBDs comprise 1.9% of the 1158-member library, consistent with the fraction of the low-fluorescence population observed via flow cytometry.

### The sequence-fitness landscape of the DNA binding domains is rugged

A fundamental question in molecular evolution is whether the emergence of new functions is mediated by the gradual partitioning of functions (smooth) or by discrete switches (rugged). On a smooth fitness landscape, we would expect a single lineage to exhibit gradual changes in activity and maintain significant promiscuity. For example, the binding specificities of proteins within the periplasmic amino acid binding protein family (same fold as the LBD of the LGF family) were shown to gradually alter over time along evolutionary lineages through the selection of promiscuous functions.^24,35^ In contrast, a rugged fitness landscape involving discrete specificity switches, would result in different functions being sparsely dispersed across the phylogeny. Our results reveal an extraordinarily rugged fitness landscape for the DBDs, in which functional repressors are dispersed within multiple lineages across the phylogenetic tree (**Fig. 1a, Fig. 2a-c**). Indeed, *LacO* repression appears to have emerged in three evolutionarily distinct lineages; the EcLacI clade, PurR/CytR clade and a third clade that we dub the functional LacI-like regulators. Notably, these clades are evolutionarily distinct and share an ancestor only at the LCA of the full LGF, which diverged ~3Gya.

**Fig. 2.**
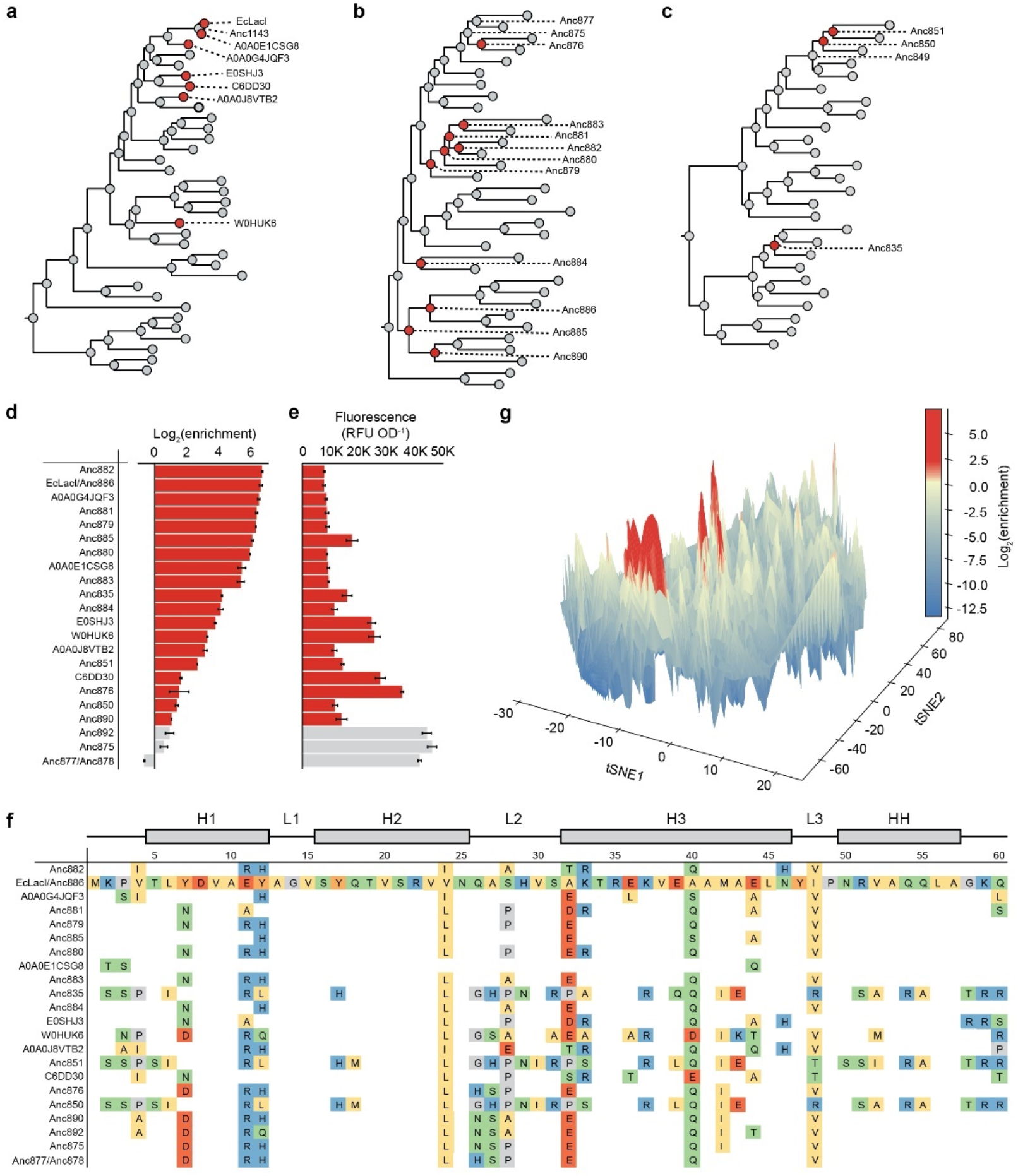
Sequence-fitness landscape and phylogenetic mapping of LacO recognition. (**a-c**), Lineages containing functional repressors of LacO. Ancestrally reconstructed sequences are numbered by tree node and extant sequences are indicated with UniProt IDs. Red nodes indicate clonally validated functional repressors of LacO within the LacI lineage (**a**), functional LacI-like regulators (**b**), and PurR/CytR-like regulators (**c**), and gray nodes indicate non-functional repressors. (**d**), Extant and ancestral DBD variants with the greatest enrichment from our high-throughput screens. Enrichment values were determined by comparing NGS distributions of the pre- and post-sorted populations. Error bars denote the standard deviation of replicate (*n*=2) sorting and NGS experiments. (**e**), Clonally assayed mean fluorescence values (*n*≥2) normalized to OD_600_. (**f**), Primary sequences of characterized ancestral and extant DBDs and overall structural topology of the DBD (top). (**g**), Sequence-fitness landscape of repressor function. Enrichment values were mapped onto the t-SNE landscape of the UniRep represented phylogenetic DBD sequences (see “methods”) to assay ruggedness. RFU, relative fluorescence units. OD, optical density.

We next traced the evolution of function within the clade in which *LacO* repression was observed, showing that recognition of LacO is sporadic within each group. For instance, in the EcLacI clade, out of the 28 ancestral sequences, only seven were repression competent. Additionally, we identify 32 instances where LacO recognition is either gained or lost between adjacent nodes (**Fig. 2a-c**). Thus, while evolutionary selection and optimization of ancestral function exist to some extent, we observe that rapid gain or loss of that same function is more dominant across the LGF phylogeny. In summary, the DBDs of the LGF family appear to be evolving across a rugged fitness landscape, leading to rapid gain/loss of function transitions.

To understand how and why sequence changes altered LacO recognition, we investigated the enrichment of different sequences, which showed a strong correlation between the enrichment ratio and the ability of a variant to repress (**Fig. 2d-f, Supplementary Fig. 6**). Clonal screening revealed a near-binary functional switch in repression competency separating enriched and depleted variants (**Fig. 2d,e**). Of the 1158-member variant library, 7 extant and 13 ancestral sequences were clonally validated as repression competent. Analysis of the sequence conservation patterns revealed among the four helices of the DBD, H2 and HH are most conserved as these are essential for LacO recognition and allosteric activation, respectively (**Fig. 2f; Supplementary Fig. 7**).^36^ Indeed, within the recognition helix (H2) residues Tyr17, Gln18 and Arg22, which directly interact with the lac operator and control DNA specificity^37^, are rarely mutated in the functionally repressive DBDs. The only exceptions include Y17H and Q18M, which have both been demonstrated to maintain lac operator binding in the genetic background of *Ec*LacI.^38,39^ Notably, ancestors 835, 850, and 851 have multiple substitutions in the HH; these variants repress GFP but are not allosterically activated by 1 mM IPTG (**Supplementary Fig. 8**), consistent with the role of HH in allosteric signaling.^36^

To obtain a global view of the sequence-function landscape, we encoded the DBD sequences as high dimensional vectors with unified representation (UniRep), a deep learning model that allows sequences with similar structural, functional, and evolutionary properties to be clustered together in space (**Fig. 2g**).^40^ This highlights the extreme ruggedness of the sequence-function landscape, with three regions that correspond to the LacI lineage, functional LacI-like regulators, and PurR/CytR-like regulators, containing most of the functional peaks. The rugged surface and isolated functional peaks suggest that there is little promiscuous binding across the family, that there is extreme epistasis with a small and finite number of functional sequences, and that there is low robustness or tolerance to mutations. In summary, our broad analysis of LacO repressor function across the 1158 sequences of this phylogenetic tree suggests that (a) LacO recognition is an easily accessible evolutionary state (appeared independently multiple times) and (b) it simultaneously exists within narrowly confined solution spaces (the fitness peaks that do exist are very small and surrounded by inactive sequences).

### Diverse substitutions shape local ruggedness among related DBDs

To better understand how the sequence space surrounding the functional repressors affects function, we identified specific mutations that affect function. This is challenging in genetically diverse datasets because neutral genetic drift and epistasis can both mask and confound sequence-function relationships. In this dataset, the context-dependence of three amino acid positions, 18, 22, and 26, illustrate how epistasis creates ruggedness (**Fig. 2a-c**). First: Gln18 in the non-functional ancestor Anc849 is mutated to Met18 (previously shown to increase affinity (2.5-fold) to LacO in EcLacI^39^) in Anc850 (alongside Glu39Leu), resulting in gain of function, followed by the introduction of Arg48Thr, Asn50Ser, and Ala52Ile in Anc851, which retains function even though these mutations are highly represented in non-functional variants (**Fig. 2c; Supplementary Fig. 9**). Thus, the deleterious effects of Arg48Thr, Asn50Ser, and Ala52Ile are mitigated by the presence of Gln18Met.^41^ Second: Arg22 is essential for LacO binding as it forms a critical base-specific hydrogen bond (**Supplementary Fig. 7b**);^42^ while it is found in every functional variant in the screen (**Fig. 2f**), 1018 non-functional variants also contain Arg22. Arg22His is among the substitutions found in Anc1143, which is the closest non-functional ancestor of EcLacI (**Fig. 2a**). Third: Asn26 in the non-functional ancestor Anc875 is mutated to His26 in Anc876 and introduces function *de novo*. However, when Asn26His is introduced alongside the Ile42Met in Anc877, there is no gain of function (**Fig. 2b**). In summary, we find subtle mutational perturbations have large effects on function, indicating a metastable state, and that epistasis contributes to the ruggedness of the landscape.

### Historical contingency and genetic drift shape evolution

While evolutionary trajectories of proteins are constrained by biological function, stochastically sampled permissive substitutions also shape the sequence landscape and contribute to historical contingency i.e., the dependence of evolutionary trajectories on historical mutations.^43–45^ From ASR alone, it is difficult to parse whether mutations are functionally essential, permissive, or neutral because ancestral and extant sequences are often separated by many substitutions. Therefore, to investigate the role of each DBD position, we used deep mutational scanning (DMS), to systematically assay all single-amino acid substitutions in EcLacI. By combining ASR with a DMS screen of EcLacI, we sought to disentangle conserved residues that have been fixed by adaptive evolution from those that arose through neutral drift.

To functionally characterize all single amino acid substitutions of the *Ec*LacI DBD (1121 variants in total), we used the same technologies (oligonucleotide chip synthesis, one-pot library cloning, FACS, and deep sequencing) as described previously for our high-throughput phylogenetic screen (**Supplementary Fig. 10**). We sorted the low fluorescence population to enrich the repression-competent variants. The number of repression-competent DMS variants (25% of the population) was 10-fold higher than from the phylogenetic library (2.5%) (**Supplementary Fig. 11a**). These were enriched to ~95% of the population after two rounds of sorting (**Supplementary Fig. 11b,c**). Sequencing of pre- and post-sorted libraries yielded enrichment ratios, normalized to native EcLacI, that were highly correlated (R^2^=0.93) between independent replicates (**Supplementary Figs. 12**)^31^.

For this DMS experiment, each DBD position has a probability distribution of the 20 canonical amino acids pre- and post-selection. We used Kullback-Lieber divergence (KLD^46^; a statistical method for comparing two probability distributions) to discern which residues are functionally restrictive (high score) and non-restrictive (low score). This showed that the restrictive and non-restrictive positions are non-uniformly distributed across the DBD, with H2 and HH largely intolerant to substitutions (**Fig. 3a, b**), consistent with the sequencing results from the phylogeny (**Fig. 2**), while the C-terminus, loop 1, loop 2 and most of H3 are robust to mutation.

**Fig. 3.**
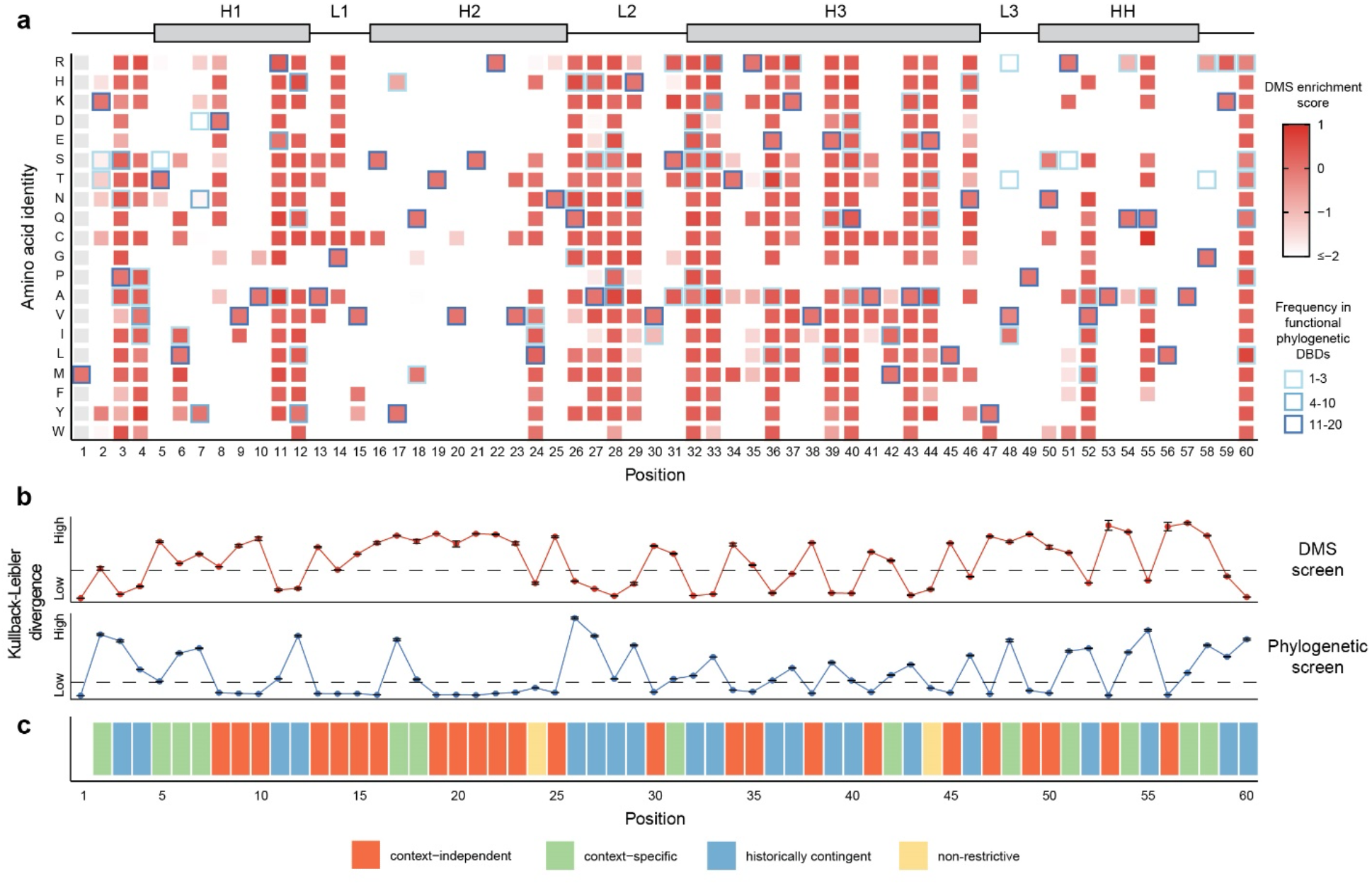
DMS to assign evolutionary roles to each position of the DBD. (**a**), Heat map showing the normalized enrichment scores (red gradient) of each EcLacI substitution after selection of repression competent DBDs. Secondary structure topology of the DBD (top) and substitution frequency (blue gradient) among functional extant and ancestral DBDs are shown. (**b**), Kullback-Liebler divergence (KLD) describes how much the amino acid probability distribution changes at each position in response to functional selection in both the DMS and phylogenetic screens. For the DMS screen, positions with high KLD are functionally restrictive, whereas low KLD positions are more tolerant to substitutions. (**c**), Assigned evolutionary roles to each DBD position based on KLD analysis of the DMS and phylogenetic screens. Context-independent positions (low phylogenetic KLD and high DMS KLD) contribute to stability and general HTH fold. Context-specific positions (high phylogenetic and DMS KLD) modulate LacO recognition. Historically contingent positions (high phylogenetic KLD and low DMS KLD) are not functionally constrained in the genetic background of *E. coli* LacI, despite sequence convergence among functional phylogenetic variants. Non-restrictive positions (low phylogenetic and DMS KLD) are generally tolerant to substitutions.

While DMS provided key insights into the functional role of each residue, it lacks an evolutionary perspective. For instance, using DMS alone we cannot distinguish between residues essential for LacO recognition (“context-specific”) and those required for stability and general fold (“context-independent”). By combining DMS with our extensive phylogenetic screen, it is possible to define an evolutionary role for each residue (**Fig. 3b**). There are four possible combinations of DMS (high/low) and phylogenetic (high/low) KLD scores, each representing a unique evolutionary role of a residue (**Fig. 3c**). Context-independent functionally restrictive residues, such as those that are structurally essential, have high DMS KLD scores (functionally restrictive) and low phylogenetic KLD scores - because positions that are conserved across all phylogenetic variants cannot undergo a shift in probability distribution upon selection and thus have low KLD scores despite potentially being functionally relevant (**Fig. 4a,b**). These context-independent residues are heavily concentrated within the core of the HTH fold and across the dimerization interface of the HH, and include essential LacO-binding residues, such as Arg22 (**Fig. 4a,b**). Context-dependent functionally restrictive positions have high KLD scores for both DMS and phylogenetic screens. These residues are critical for LacO recognition, either making direct contact with the operator DNA in both the major and minor grooves, helping to orient the DBD for LacO binding, or interacting with the LBD for allosteric communication. Context-dependent residues include Tyr7, Tyr17 and Gln18 and Met42 – all of which contact LacO or are associated with an epistatic change-of-function, such as in Anc877 (**Fig. 4c,d**). Historically contingent positions have low DMS and high phylogenetic KLD scores i.e., while mutationally robust in terms of EcLacI function, they are restricted in phylogenetic analysis owning to their deterministic role in evolution (**Fig. 4e,f**). Many of these residues are located adjacent to context-dependent functionally restrictive positions, consistent with a role in modulating their effect, such as Asn26 (historically contingent) and Met42 (context-specific) (**Fig 4f**). Functionally non-restrictive positions have low KLD scores for both DMS and phylogenetic screens. Only two solvent-exposed residues, Val24 and Glu44, fall within this category. Altogether, this analysis demonstrates that the evolutionary role of all residues within the DBD of EcLacI can be revealed through a statistical analysis of DMS and ASR data.

**Fig. 4.**
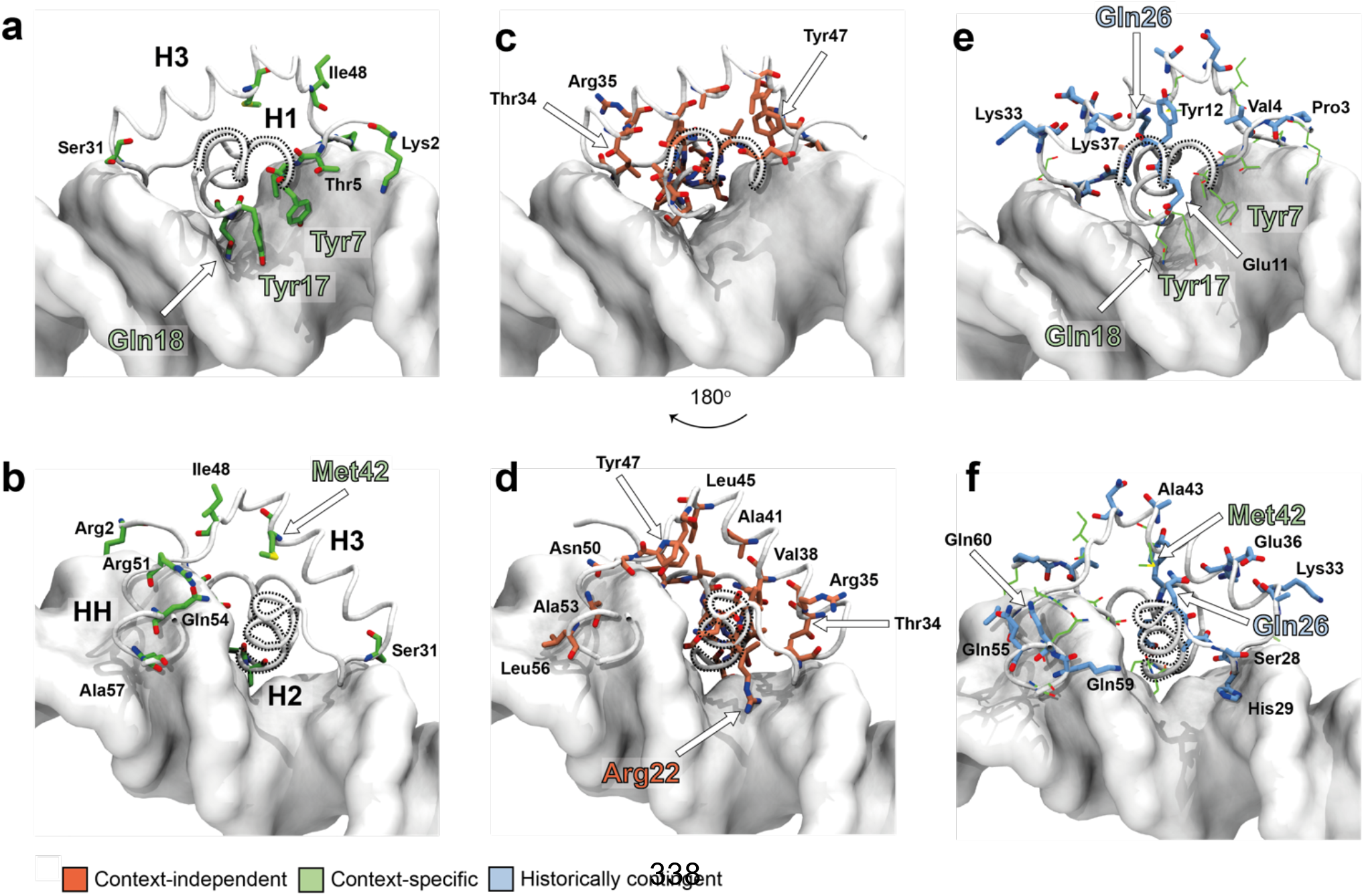
Structural role of DBD mutations. (a) Context-specific mutations in the LacO major groove, where Tyr7, Tyr17 and Gln18 make essential contacts with the DNA. (b) context-specific mutations in the minor groove. Context-specific mutations are dominated by locations that contact the DNA, particularly in the major groove. (c) Context-independent mutations in the major groove. Context-independent mutations cluster around the core packing of the DBD. (d) Context-independent mutations in the minor groove. Context-independent mutations also include those essential for DNA-recognition, such as Arg22. (e) Historically contingent mutations in the major groove. Context specific mutations are represented as lines (green), historically contingent are represented as sticks (cyan). (f) Historically contingent mutations in the minor groove. Historically contingent mutations cluster around the context specific mutations that they set the background for. For example, Met42 and Gln26 (epistatically linked) are oriented towards one another on H3 and H1, respectively.

### Operator asymmetry contributes to the rugged fitness landscape

Despite asymmetric DNA operator sequences having lower affinity for DBDs than symmetrical sequences,^47^ they are overwhelmingly dominant throughout evolution and must therefore confer some selective benefit. To investigate this, we performed *in vitro* DNA-binding assays using surface plasmon resonance (SPR; **Fig. 5**). We find that all functional variants tested (Anc880, Anc881 and Anc882) bind the symmetrical LacO_sym_ with *K*_d_ constants greater than an order of magnitude lower than their asymmetric operator counterparts (LacO and LacO_1_), confirming that operator asymmetry significantly reduces a repressor’s DNA-binding capacity in ancestral, as well as extant DBDs (**Fig. 5a-c**; **Supplementary Fig. 13**).

**Fig. 5.**
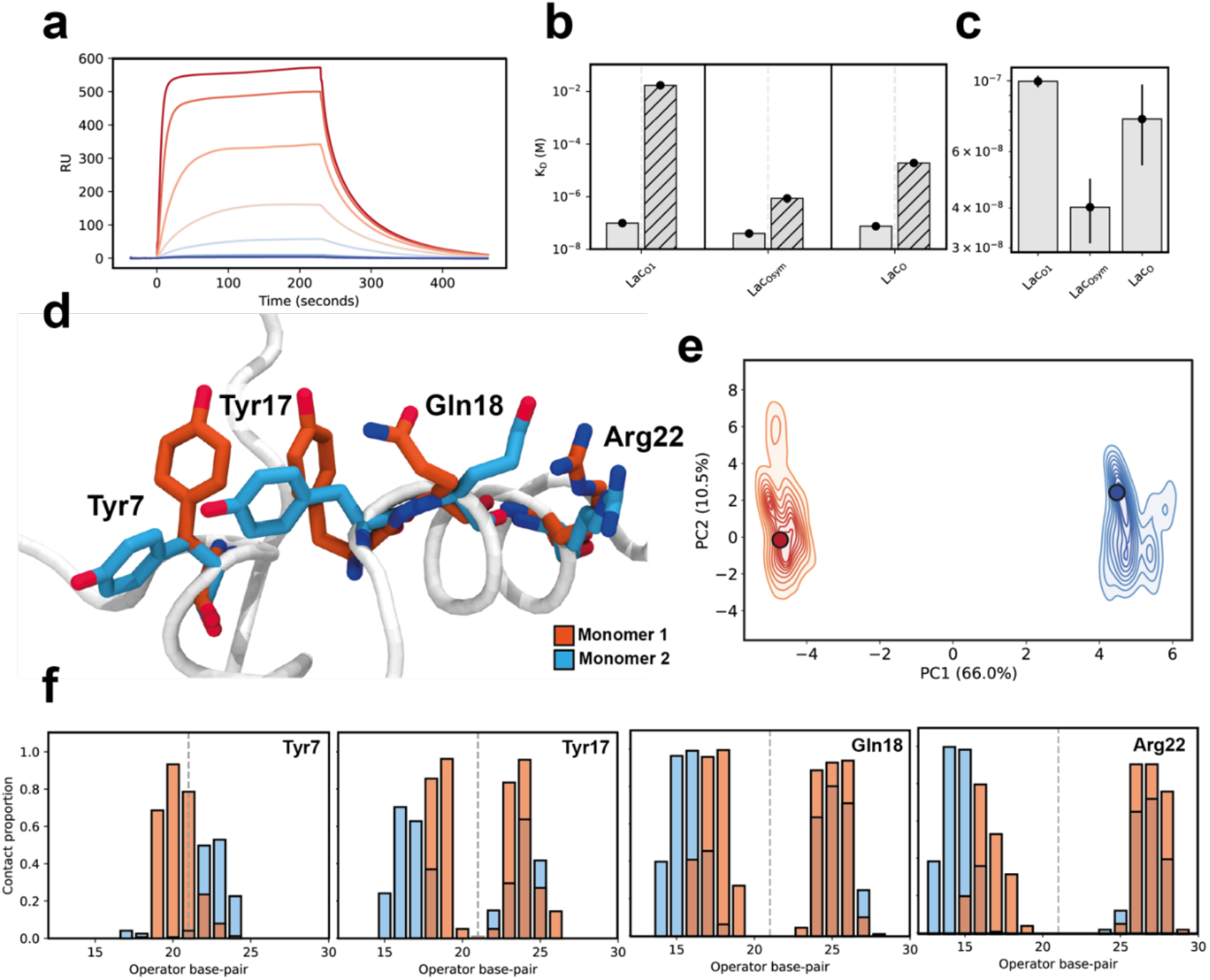
Transcriptional regulator asymmetry and fitness landscape ruggedness. (a) Multi-cycle kinetics SPR sensogram for Anc882 LacO binding. (b) Anc882 *K_d_* values, determined by SPR for LacO_1_, LacO_sym_ and LacO DNA binding when repressed (solid) and induced with 10 mM IPTG (hashed). (c) Magnified view of repressed *K*_d_ values. (d) Conformational snapshots of the most sampled conformations for monomers 1 (red) and 2 (cyan). Dominant conformations of Tyr7, Tyr17, Gln18 and Arg22 are shown from principal component analysis (PCA; panel e). (e) PCA of Anc882, showing monomers A (red) and B (blue). Conformational snapshots presented in (d) are shown as circles. (f) Distribution of key protein-DNA contacts in Anc882 and LacO. DNA residues (X-axis) have been aligned for visualization.

To investigate the molecular basis for binding of asymmetrical operator sequences, we performed all-atom molecular dynamics (MD) simulations on models of Anc880 and 882, as well as EcLacI, complexed with LacO (**Fig. 5**). Each system was simulated with 10 independent 60 ns replicates (600 ns total sampling time) (**Supplementary Fig. 14**). Previous studies have identified motions related to DNA-binding specificity in *Ec*LacI occur over ns – μs timescales, consistent with the timescale of our simulations.^48,49^ We observe asymmetry in the protein-DNA interactions between the two monomers that constitute the functional dimer (**Fig. 5d,e**). Although both monomers interact with the DNA via the same residues, the relative contributions these residues make to the overall binding energy and their DNA contacts are distinct. In Anc882, which is the variant with the greatest LacO affinity, we identified Tyr7, Tyr17, Gln18 and Arg22 as being essential for DNA recognition and binding. However, the conformations of each of these residues differ between monomers to accommodate the respective operator half-sites and principal component analysis (PCA) reveals that backbone motions differ between either monomer (**Fig. 5d-f**). Identical analyses of apo-Anc882 trajectories found that in the absence of DNA, backbone motions are nearly indistinguishable between monomers (**Supplementary Fig.14**). This analysis is consistent with NMR structures of the EcLacI DBD complexed with the natural LacO_1_ operator^50^, and alternate DNA binding modes have been alluded to by analyses of LGF repressors and EcLacI mutants.^51^ Thus, asymmetrical operator sequences impose a complex selective pressure: the DBDs must be able to undergo a conformational change to bind two distinct half-sequences with physiologically relevant affinity. Being subject to multiple selection pressures associated with either half-site to maintain complete operator binding is consistent with the extreme ruggedness we observe (**Fig. 2**).

The demand on the DBD to simultaneously evolve high affinity for two distinct half-site DNA sequences using a single binding site (as well as allosteric regulation with the LBD) means that the fitness landscape we observe across this phylogeny can be viewed as a composite of two or more fitness landscapes. To test this, we performed *in silico* evolutionary simulations on small model systems using theoretical elementary landscapes as a model (**Fig. 6**)^52–54^. Elementary landscapes derive from a robust and well-established graph-theoretic approach to combinatorial landscape optimization problems^54^; specifically, they are the orthogonal eigenvectors of the graph-Laplacian of a sequence space graph. For our purposes, they provide a theoretically rigorous way of testing hypotheses that pertain to the ruggedness of a fitness landscape. Using a linear combination of elementary landscapes, each representing a different hypothetical fitness function, we find that the combination of two fitness landscapes can indeed be more rugged than either constitutive fitness landscape is alone, akin to constructive/deconstructive wave interference (**Fig. 6**). This demonstrates that ruggedness can emerge from the composition of otherwise smooth fitness landscapes and predicts that proteins with a single function, such as enzymes or solute binding proteins, may evolve over smoother fitness landscapes than proteins such as LGF repressors, where multiple and complex fitness constraints are imposed simultaneously.

**Fig 6.**
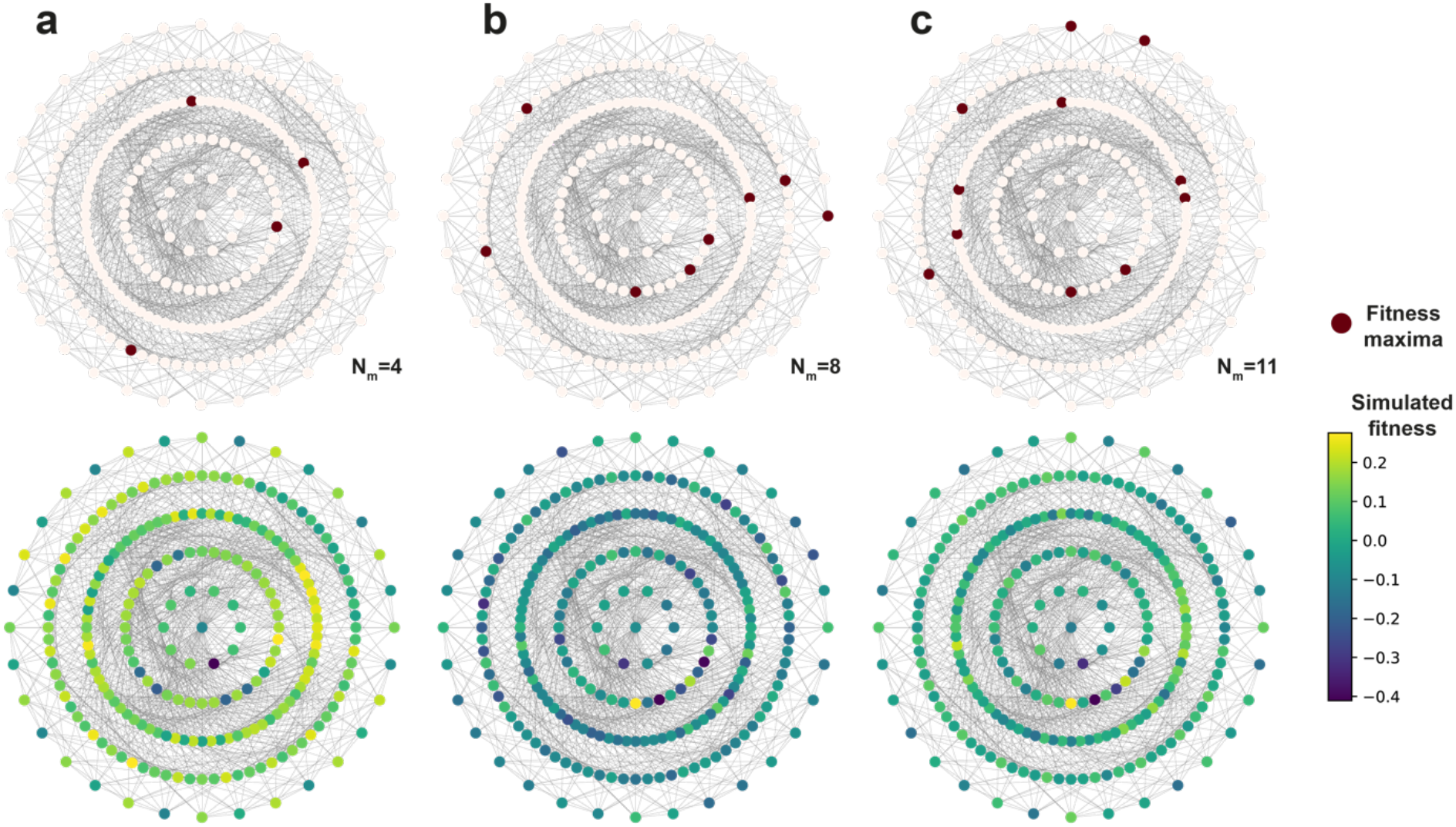
Elementary landscape model simulations. 2^nd^ and 3^rd^ elementary landscapes in panels (a), (b) respectively. Each vertex represents a unique genotype, colored fitness maxima (top) and quantitative simulated fitness (bottom). Edges connect genotypes accessible by a single mutation. (a) and (b) have 4 and 8 fitness maxima (N_m_), respectively (where N_m_ is the number of fitness peaks where all neighboring genotypes have a lower fitness). (c) A linear combination of 2^nd^ and 3^rd^ elementary landscapes. The composite landscape is characterized by 11 fitness maxima, more than either constituent landscape, and is more rugged.

## DISCUSSION

These results show that the fitness landscape for operator binding by DBDs of the LGF is extremely rugged i.e., the viable fitness space for LacO recognition and repression is narrow and highly localized. Interestingly, this ruggedness has long been alluded to by mutational studies, but never comprehensively observed or studied prior to our work.^5^ In contrast, most of the protein families studied by ASR to date demonstrate protein evolution over smooth fitness landscapes where gradual changes in function can be observed.^24,35,45,55–58^ Even those characterized by extensive epistasis have well-defined mutational trajectories from semi- or non-functional intermediate sequences to functional ones^44,59^. The profound difference in evolutionary dynamics between these studies and the LGF may reflect fundamentally different biological and physical constraints. While any complex molecular trait can be described as a combination of arbitrarily smooth fitness functions (i.e. folding, substrate-recognition, electrostatic preorganization, immunogenicity, among many catalytic and structural parameters^60^) DNA recognition among LGF regulators is particularly rugged due to the compounding of two DNA-binding landscapes that likely have complexity in isolation; if either half-site requires the correct folding, dynamical ensemble, electrostatic and shape complementarity (among likely many other traits), a protein sequence that binds both half-sites must compound the molecular requirements for both of the half-sites.

One notable aspect of the rugged DBD landscape is the evolutionary metastability of repression: descendants of a repression competent ancestor are seldom functional with the same operator sequence and mutations have extreme epistatic relationships. This indicates that both selection pressure for LacO specificity has been variable and that LacO specificity has emerged independently numerous times throughout evolutionary history, each time in different sequence contexts. Like the notion of an epistatic ratchet, where a phenotype becomes completely inaccessible within a specific background^43,61^, the metastability we observe occurs through non-specific mutations that counteract or reciprocate the fitness effects of previously fixed mutations in backgrounds that are on the edge of a binary phenotype, making a few genetic changes sufficient to rewire specificity. Using a combination of ASR and DMS, we have comprehensively mapped historical contingency within the DBD showing that permissive sites, which are essential in an evolutionary context and underlie the epistasis and metastability, can be identified.

The rugged landscape, the asymmetry of the DBD:DNA complex and the evolutionary metastability of the repressor function are all interesting observations, but how do they relate to the physiological role of these proteins? Since effective genetic regulation is paramount to organismal fitness, precise and binary interactions are essential. To impart a selective advantage, regulators must be highly specific for their cognate operator sequences. Unlike promiscuous enzymatic activity, promiscuous DNA-binding causes metabolic cross-regulation and severe disruption of cellular function.^62,63^ Thus, we hypothesize that this ruggedness fitness landscape is an intrinsic aspect of fitness for the LGF: asymmetric operator half-sites have evolved as a mechanism to minimize metabolic crosstalk between regulators and create an evolutionary dynamic in which activity is essentially binary. Over a rugged fitness landscape, as we observe, DNA-binding specificity is metastable, with rapid gain/loss of function, essentially eliminating the risk of non-specific DNA-binding and metabolic cross-regulation between diverging regulators. This suggests that inherently epistatic protein folds, such as that of the DBD, have likely been evolutionarily selected for regulatory purposes. Indeed, previous ASR studies have shown that epistasis is a hallmark of several classes of eukaryotic transcription factors,^59,64–67^ and recent results have highlighted the importance of molecular unpredictability in transcription factor divergence.^68^ Interestingly, similar observations of rapid *de novo* promoter emergence have also recently been made, indicating that the evolutionary dynamics we observe in regulatory proteins may also be present in their DNA binding partners.^69^

Altogether, our analysis of the sequence-function landscape of the LGF DBDs has led to several valuable insights. Most importantly, we prove a molecular level explanation for the high fidelity that has long been observed in gene regulation by proteins of the LGF family, showing that it is an intrinsic property of the rugged fitness landscape that is itself a function of the biophysical properties of DBD structure and asymmetry of DNA operator sequences. Future work could investigate whether rugged fitness landscapes, and functional partitioning through asymmetry, are common when the physiological function of the protein family makes promiscuous activity deleterious, as for these transcriptional regulators.

## METHODS

### Phylogenetic inference

To collate an expansive dataset of the full LGF, EcLacI, RbsR, MalR, SacR/ScrR, GalR, GalS, AscG, PurR, CytR, and CcpA were each individually queried against the NCBI non-redundant protein database using pBLAST. ~250 sequences were retrieved from each individual BLAST search, using an e-value significance threshold of 1.0E-10. Redundancy in this dataset was removed to 90% sequence identity using CD-HIT^70^ and incomplete sequences, or sequences with poorly conserved (<1% of sequences) indels were manually removed. Sequences were aligned using the ESPRESSO protocol of T-COFFEE. ^71^ This alignment was manually refined to remove poorly conserved sequences or insertions and benchmarked against available LGF X-ray crystal structures. Phylogenetic inference was performed in IQ-TREE.^72^ The ML model, LG+R9,^73,74^ was fitted using ModelFinder (as implemented in IQ-TREE).^74^ Tree-search was performed using default parameters and branch supports were computed as ultra-fast bootstrap approximations.^75^ Tree search was repeated for 10 independent replicates, which were tested for statistical equivalency by the approximately unbiased (AU) test conducted to 10000 replicates.^33^ An additional inference was performed using the same protocol, however, only including the N-terminal 60 columns of the LGF dataset to determine how much evolutionary signal the DBD provides to the full-length sequences (Supplementary Fig 2). Ancestral sequences were reconstructed on the single topology presented in Fig. 1 using CodeML from the PAML software package.^76^ The sequence evolution model for ASR was manually set to LG+G4^73^ and indel events were processed in the ancestral sequences according to the principle of parsimony.

### Construction of GFP reporter strain

The *E. coli* strain DH10β was modified by lambda Red recombineering to insert a superfolder GFP gene driven by the pLlacO^77^ promoter and a kanamycin resistance cassette at the LacI locus to generate the GFP “reporter” cell line. The temperature sensitive plasmid pKD46 (Genbank AY048746) was used for arabinose-inducible expression of Red recombinase.^78^ Linear double-stranded donor DNA was amplified off a pSC101 plasmid using overhang primers to add 50bp homology arms directed to the 5’ end of the LacI gene. A frozen glycerol stock of DH10β transformed with the pKD46 plasmid was struck out on LB carbenicillin (100μg/mL) and incubated at 30°C overnight. A colony was selected and inoculated into 5mL LB carbenicillin and grown for 16h in a shaking incubator at 30°C. Cells were diluted 50X into 25mL of LB carbenicillin and grown to an OD of 0.1. Red recombinase was induced with 100mM L-arabinose and cells were grown to an OD of 0.6. The cells were harvested and prepared for electroporation. 25μL of cells were transformed with 500ng of donor DNA. Cells were recovered for 2h in SOC media in a shaking incubator at 37°C and plated on LB kanamycin (30 μg/mL). The transformants were incubated at 37°C overnight to cure the cells of the temperature sensitive pKD46 plasmid. A visibly fluorescent colony was selected and inoculated into 5mL LB kanamycin to be grown for 16h in a shaking incubator at 37°C. The genome modification was confirmed via sequencing and a glycerol stock was stored at −80°C.

### General library DNA assembly

Plasmids were constructed using standard molecular biology techniques of PCR and Golden Gate assembly with Kapa HiFi DNA Polymerase (KAPA Biosystems), restriction enzymes (NEB), T4 DNA Ligase (NEB), and Antarctic Phosphatase (NEB).

Oligonucleotides encoding residues 2-60 of all LGF DBDs (1158 variants in total) or all single amino acid substitutions of the *E. coli* LacI DBD (1121 variants in total) were synthesized as single-stranded oligonucleotide pools (Agilent Technologies). Extant DBDs exceeding 60aa long were truncated from the N-terminus to 60 residues in length to comply with DNA synthesis restrictions. Oligonucleotides were converted to double-stranded DNA using 15 cycles of PCR amplification and purified on a spin column (EZNA Cycle Pure kit from Omega BioTek). A pSC101 backbone containing extant *E. coli* LacI gene under control of the strong pLtetO promoter and a spectinomycin resistance gene was amplified using a primer pair encoding BsaI cut sites that matched the DBD insertion location of both oligonucleotide libraries. The amplified backbone was treated with Dpn1 for 2h at 37°C and purified using a spin column. The backbone was treated with BsaI-HF v2 for 2h at 37°C, Antarctic phosphatase for 1h at 37°C, and subsequently purified using a spin column. A Golden Gate assembly reaction (30 cycles of 37°C for 5min and 16°C for 5min) was performed using 0.042 pmol pSC101 backbone and 0.21 pmol amplified DBD library in a 1:5 molar ratio. The assembled product was dialyzed on a semi-permeable membrane (Millipore) for 1h at 25°C against dH_2_O. A 25μL aliquot of electrocompetent DH10β *E. coli* cells were transformed with 2μL of the dialyzed product using electroporation. Cells were recovered for 1h in SOC media in a shaking incubator at 37°C and dilutions were plated on LB Spectinomycin (50 μg/mL) to calculate transformation efficiency (>10^6^ CFU/mL). Remaining recovered cells were diluted 5×, incubated for 6h, and then diluted 50X for overnight growth in a shaking at 37°C for 16h. Library plasmids were extracted using a DNA miniprep kit and 1μL (~100ng) was used to transform 25μL of electrocompetent reporter cells following the transformation protocol described above. Glycerol stocks of the reporter cells transformed by the preselected phylogenetic and DMS plasmid libraries were stored at −80°C.

### Fluorescence-activated cell sorting

Thawed glycerol stocks of reporter cells containing the LGF DBD or DMS libraries were used to inoculate (50μL each) 5mL of LB kanamycin (30μg/mL) / spectinomycin (50μg/mL) in duplicate. Cells were grown in a shaking incubator at 37°C for 16h and subsequently diluted 50X in PBS (137mM NaCl, 2.7mM KCl, 10mM Na2HPO4, 1.8mM KH2PO4) for sorting. Sorting was performed with a Sony MA900 cell sorter. Cells were excited with a 488nm laser and GFP fluorescence was monitored through a 525/50 filter. A sorting gate was drawn to isolate the low-fluorescence populations of the LGF DBD (2.5-2.6% of total population, 250,000 cells collected) and DMS (25.4-25.6% of total population, 500,000 cells collected) libraries. Cells were sorted into 1mL of LB, and the total volume was adjusted to 5mL after sorting. Cells recovered for 1h at 37°C in a shaking incubator before antibiotics were added (kanamycin and spectinomycin), and incubation was continued for another 15h. Glycerol stocks were made and stored at −80°C. Plasmids were harvested using a DNA miniprep kit and 1μL (~100ng) was used to transform 25μL of fresh electrocompetent reporter cells. The procedure described above was repeated for a second round of low-fluorescence sorting using the same sorting gates to further enrich for repression competent variants.

### Next-generation sequencing

We used deep sequencing to evaluate presorted and sorted LGF and DMS populations. The DBD was amplified in two stages with Kapa HiFi DNA Polymerase (KAPA Biosystems) for tailed amplicon sequencing. The first PCR reaction was performed with overhang primers that anneal to 5’ and 3’ constant regions surrounding the DBD. The overhangs add a variable N region (to enhance nucleotide diversity), and a portion of the universal Illumina adapter. The first reaction was performed using 14 cycles with 1ng of extracted plasmid DNA (DMS or LGF library, presorted or sorted) as template. The product was purified on a spin column (EZNA Cycle Pure kit from Omega BioTek). A second PCR reaction was used to add the index (for pooled sequencing) and the Illumina ‘stem’. This amplification was performed using 10 cycles with 10ng of the purified product from the first reaction as template. The amplicons were purified and used for deep sequencing. Samples were sequenced on an Illumina MiSeq System, using a MiSeq Reagent Kit v2 (2×250 cycles) following the manufacturer’s documentation.

Paired-end Illumina sequencing reads were merged with PEAR (Paired-end read merger).^79^ Phred scores (Q scores) were used for quality filtering. Reads with an expected number of errors exceeding 1 were removed and total read counts for each DBD variant were computed (**Supplementary Figs. 3,10**). Raw read counts were highly correlated (R^2^≥ 0.97) for all replicate samples. Enrichment for each variant was computed using

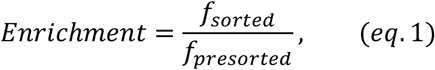

where *f_presorted_* and *f_sorted_* are relative read count frequencies before and after enrichment of repression competent variants, respectively. Higher enrichment indicates tighter repression of GFP and higher affinity to LacO. For the DMS library, enrichment scores (Escore) were computed by normalizing enrichment to WT and applying a log2 transformation. Thus, the WT DMS Escore is set to 0, Escore > 0 indicates improved function, and Escore < 0 indicates reduced function.

Kullback-Liebler divergence of a specific position is given by the following:

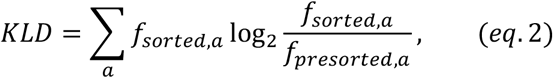

where the sum is over all 20 canonical amino acid identities, and *f_sorted,a_* and *f_presorted,a_* are the relative frequencies of an amino acid *a* in the repressed sorted or presorted distributions, respectively. If the relative frequency of an amino acid at a position is zero, then the value was excluded from the summation.

### Microplate fluorescence assay for clonal characterization

The presorted and repressed sorted LGF DBD libraries were struck out on LB kanamycin (30μg/mL) / spectinomycin (50μg/mL) plates. Colonies were selected and inoculated into 150μL LB kanamycin/spectinomycin in a 96-well plate. Cells were incubated at 37°C in a microplate shaker for 8h. The cultures were diluted 50X in fresh media in a 96-well plate with varying concentrations of IPTG (0, 0.1, 0.5, 1, 5, 25, 100 and 1000μM). Diluted cultures were incubated at 37°C in a microplate shaker for 14h. GFP fluorescence and OD_600_ were measured in a BioTek Synergy HTX Multi-Mode 96-well plate reader. Fluorescence was normalized to OD to account for differences in cell density. Assayed colonies were sequenced to identify the DBD variant. The mean and standard deviation of normalized fluorescence for replicates (n ≥ 2) for each concentration of IPTG were used to fit these dose-response curves to the Hill–Langmuir equation:

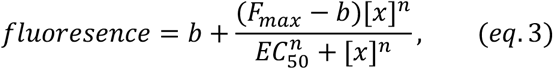

where *b* is the basal fluorescence (measure of LacO affinity in the absence of inducer), *F_max_* is the maximum fluorescence signal achieved at 1mM IPTG, *EC*_50_ is the half maximal effective concentration, *x* is the concentration of IPTG, and *n* is the Hill coefficient.

### Protein expression and purification

Anc880, 881 and 882 were synthesized and cloned into the NdeI/XhoI multiple cloning site of pET-28a(+) with a C-terminal 6× histidine tag. All three ancestral proteins were recombinantly expressed in BL21(DE3) lab strain *E. coli* at 25 °C for 16 hours in Luria-Bertani (LB) media. Protein expression was induced with 10 mM IPTG once media OD_600_ reached approximately 1.5 AU. Cells were pelleted by centrifugation at 5,000 RPM for 15 minutes and were lysed by sonication once resuspended in buffer A [20 mM PBS, 150 mM NaCl, 20 mM imidazole (pH 7.4)] containing the recommended amount of *Serratia marscecens* Turbonuclease (Sigma-Aldrich). The lysate was pelleted by centrifugation at 12,000 RPM, 4 °C for 1 hour before being filtered (0.22 μM) and loaded onto a prepacked 5 mL HisTrap FF immobilized metal ion affinity chromatography column (GE lifescience) equilibrated in buffer A. The loaded column was washed for 5 column volumes in buffer A and 3 column volumes in 94% buffer A/ 6% buffer B [20 mM PBS, 150 mM NaCl, 250 mM imidazole (pH 7.4)], before being eluted with 100% buffer B on an AKTA Start FPLC instrument (GE lifescience). IMAC purified protein was then filtered (0.22 μM) and purified by size exclusion chromatography (SEC) on a prepacked HiLoad superdex 200 16/60 SEC column (GE lifescience) equilibrated in SEC buffer [20 mM PBS, 150 mM NaCl (pH 7.4)] and run on an AKTA FPLC system (GE lifescience).

### Surface Plasmon resonance

Proteins purified to homogeneity by IMAC and SEC were buffer exchanged into SPR running buffer [10 mM HEPES, 300 mM NaCl, 3 mM EDTA, 0.05% (v/v) tween-20 (pH 7.4)] and diluted into a log2 dilution series from 250 nM - 3.91 nM for repressed binding samples and 10 μM – 78.1 nM for samples induced with 70 μM IPTG. _Double_-stranded DNA oligonucleotides for LacO, (TGTGTGGAATTGTTATCCGCTCACAATTTCACACA) LacO_1_ (TGTGTGGAATTGTGAGCGGATAACAATTTCACACA), LacO_sym_ (TGTGTGGAATTGTGAGCGCTCACAATTTCACACA), and random DNA (AGGTCAAAAAGCCAGTGGTTATTTTAAGATGTCGC) were synthesized and 5’-biotinylated (IDT). All SPR experiments were performed on a Biacore 8K instrument (GE lifescience). An SA streptavidin Biacore chip (GE lifescience) was activated with 1 M NaCl and 20 mM NaOH before four channels were charged with approximately 400 RU of each respective DNA ligand. Binding affinity was determined through multi-cycle kinetic analysis at the 7 serial log_2_ protein dilutions, and one buffer only blank. All protein analytes were run at 50 μL/min for a ligand contact time of 120 seconds. After each analyte cycle, the ligand was regenerated with 3 M NaCl run at 20 μL/min for a contact time of 60 seconds. The K_d_ of DNA binding was determined from a multi-cycle model fitted to the reference subtracted sensorgrams in the Biacore 8K evaluation software.

### Molecular dynamics simulations

Models for Anc880 and Anc882 were generated from X-ray crystallography coordinates of EcLacI bound to LacO_sym_ (PDB: 1EFA)^80^ using FoldX^81^. The LacO_sym_ DNA ligand in 1EFA was mutated to the LacO sequence that was experimentally tested using X3DNA. ^82^ To avoid DNA terminal fraying, 5’ and 3’ termini were extended with sequences GGAAT and TCC respectively in X3DNA for all models (Full DNA sequence: GGAATTGTTATCCGCTCACAATTCC). Simulations were performed in the GROMACS MD engine using the Amber ff14SB^83^ force field with PARMBSC1^84^ nucleic acid parameter set. MD integration used a timestep of 2 fs. Columb interactions were modelled using particle mesh Ewald (PME) with a cut-off radius of 1.0 nm. Leonard-Jones interactions used a cut-off scheme with a radius of 1.0 nm. Hydrogen bonds were constrained using the LINCS algorithm. ^85^ For each system, 10 independent replicates of equilibration (from the NVT ensemble onwards) and production MD were performed. Each protein-DNA complex was put in a rhombic dodecahedral box extending at least 1.0 nm from the closest atom to the box boundary in all directions. Simulation boxes were filled with TIP3P water molecules, and the charge was neutralized with sodium and chloride ions. To replicate physiological conditions, each box was additionally filled with 150 mM NaCl. All systems were energy minimized by steepest descent with a step size of 0.01 nm until the greatest force was less than 1000 KJ/mol/nm. The systems were equilibrated to 300 K in the NVT ensemble with harmonic restraints on all heavy atoms for 100 ps using a velocity rescale thermostat and brought to 1 Bar pressure in the NPT ensemble for 100 ps using a Berendsen barostat. Following temperature and pressure equilibration, unrestrained production MD was performed for 100 ns in the NPT ensemble using a velocity rescale thermostat and a Parrinello-Rahman barostat with references of 300 K and 1 Bar, respectively. Following periodic boundary corrections, all trajectories were analyzed in MDTraj and every 10^th^ frame was sampled. All production MD was performed on the Australian National Computing Infrastructure GADI supercomputer.

### Elementary landscape simulations

Elementary landscapes were calculated via eigendecomposition of the graph adjacency matrix corresponding to a combinatorial sequence space over an alphabet of 3 amino acids and 5 number of positions. Elementary landscapes were indexed by order (e.g. first-order, second-order and so on), and composition of landscapes performed combinatorically between 1st and 2nd order, 1st and 3rd order, and 1st and 3rd order landscapes. The composition operation was defined as a simple linear combination of landscapes (in analogy to superposition of waves in physics). All graph construction and matrix operations were performed in NumPy and Networkx. The number of local maxima were calculated using custom functions. Graph plotting was performed in Networkx using custom functions.

## ACKNOWLEDGEMENTS

This work was supported in part by the ARC Centre of Excellence in Synthetic Biology (CE200100029), the ARC Centre of Excellence in Peptide and Protein Science (CE200100012), the Great Lakes Bioenergy Research Center, U. S. Department of Energy, Office of Science, Office of Biological and Environmental Research under Award Number DE-SC0018409 (S.R and A.T.M) and the NIH Director’s New Innovator Award DP2GM132682 (S.R). The content is solely the responsibility of the authors and does not necessarily represent the official views of the NIH, Department of Defense, Department of Energy, or other federal agencies.

## AUTHOR CONTRIBUTIONS

A.T.M., M.A.S., C.J.J, and S.R. conceptualized the project. M.A.S. performed the phylogenetic analysis and ancestral protein reconstruction. A.T.M. created designs, performed sorting and NGS experiments, and Rosetta stability calculations. M.A.S. performed binding experiments and operator half-site analyses. M.S. simulated the NK model landscapes. A.T.M., M.A.S., C.J.J, and S.R. drafted the manuscript, A.T.M., and M.A.S. created the figures. C.J.J. and S.R. supervised the work. All authors edited and approved the final manuscript.

---

**Supplementary Fig. 1.**
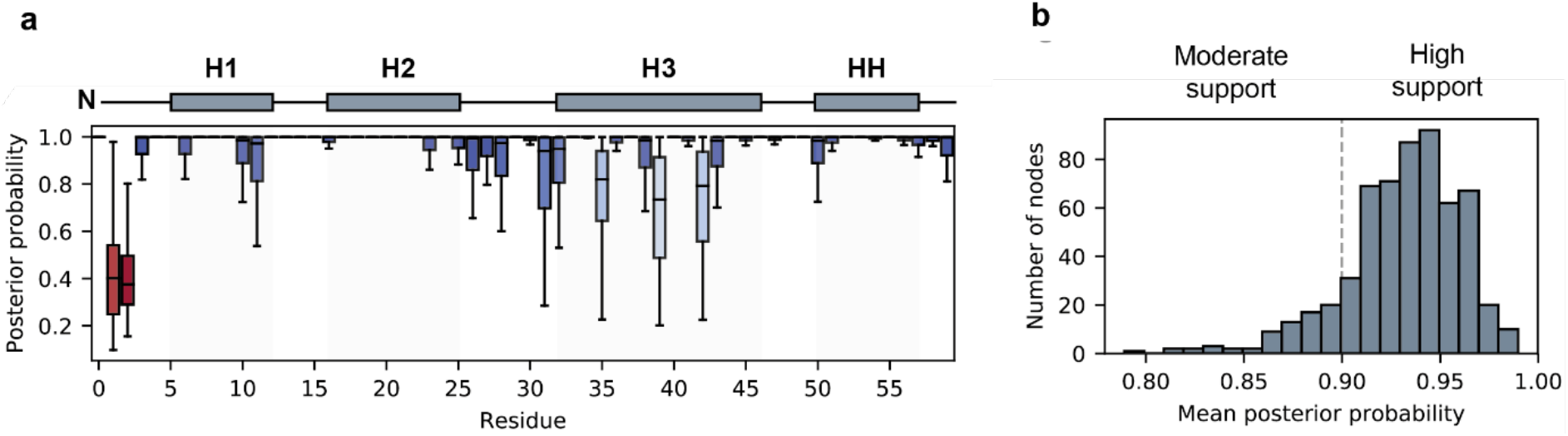
Ancestral sequence reconstruction of prokaryotic transcriptional regulators. (**a**), Site-wise posterior probability distribution averaged over all reconstructed ancestral sequences. (**b**), Distribution of posterior probabilities averaged over the DBD sequence. Most ancestral sequences are reconstructed with high posterior probability (>0.9 mean posterior probability) and very few are reconstructed with less than moderate (<0.8 mean posterior probability). Helix 2 and the hinge-helix, which are most important for DNA-recognition, are reconstructed unambiguously.

**Supplementary Fig. 2.**
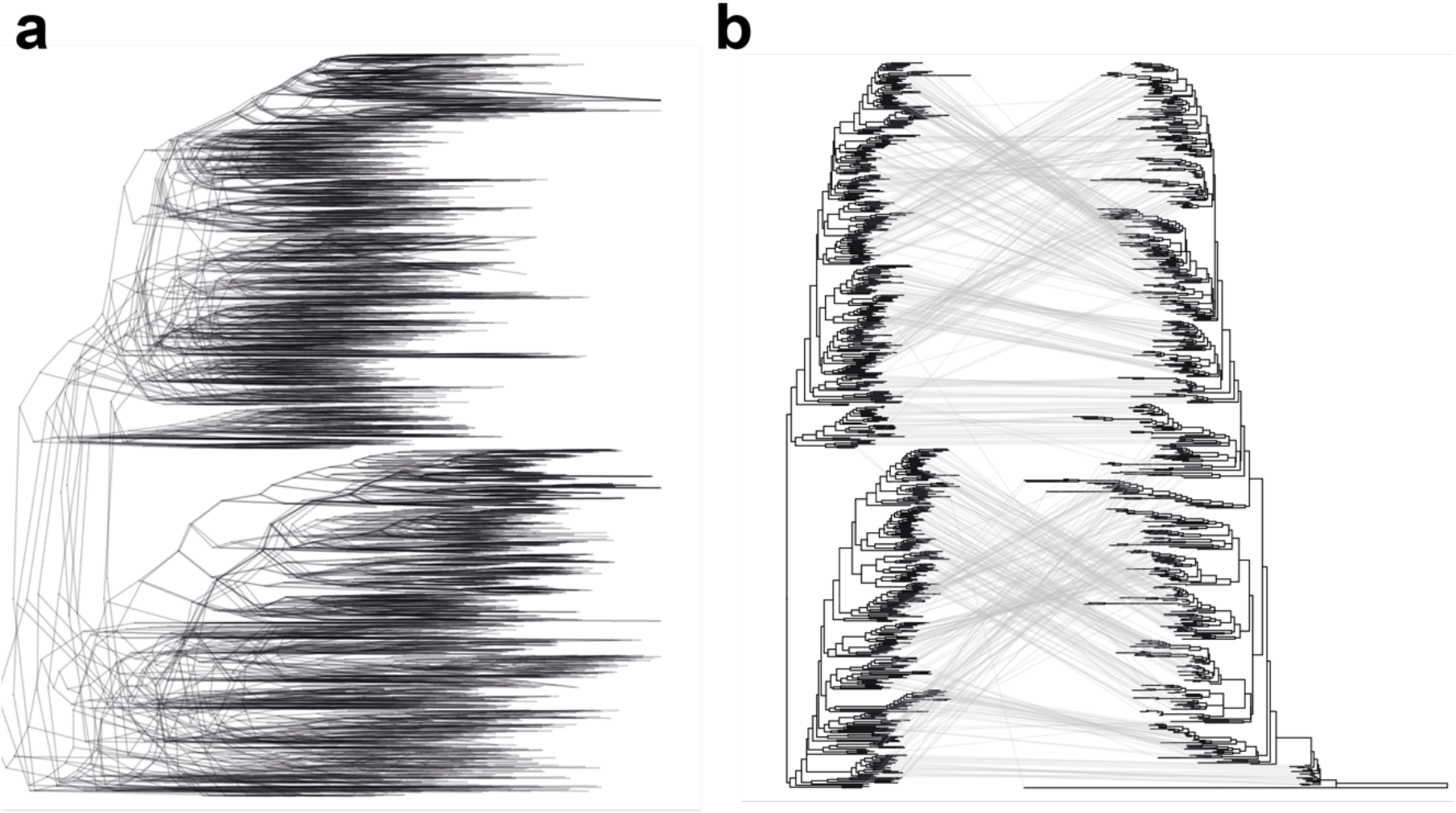
Tree topology testing. (**a**), All 10 reconstructed topologies that passed the AU test conducted to 10000 replicates. Each independent tree search converges on a similar topology. (**b**), Comparison the full DBD and LBD topology presented in main text Fig. 1 (left) and a tree-search performed under the same criteria using only the DBD. Most of the phylogenetic signal in the inference comes from the large LBD sequence, as the DBD-only topology is incongruent with the full-length topology.

**Supplementary Fig. 3.**
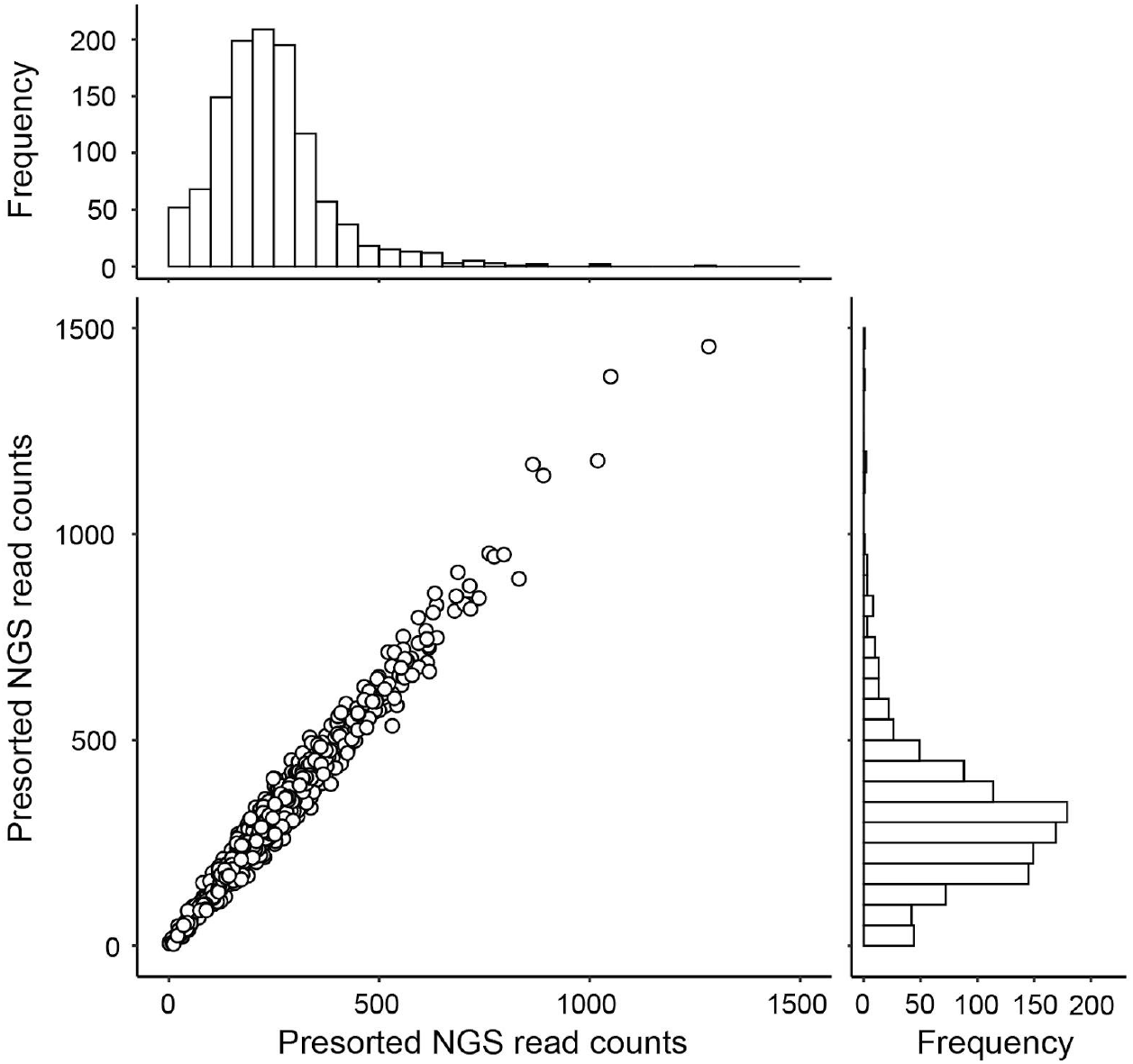
Presorted library distribution of extant and ancestral DBD variants. Raw read counts of all 1158 DBD variants from replicate next-generation sequencing runs (*n*=2). All variants were present in the pooled library (median=250±40 reads) prior to selection of functional repressors. Histograms of presorted read counts show that DBD variants were approximately normally distributed with a moderate right-shift (skewness=1.6±0.1).

**Supplementary Fig. 4.**
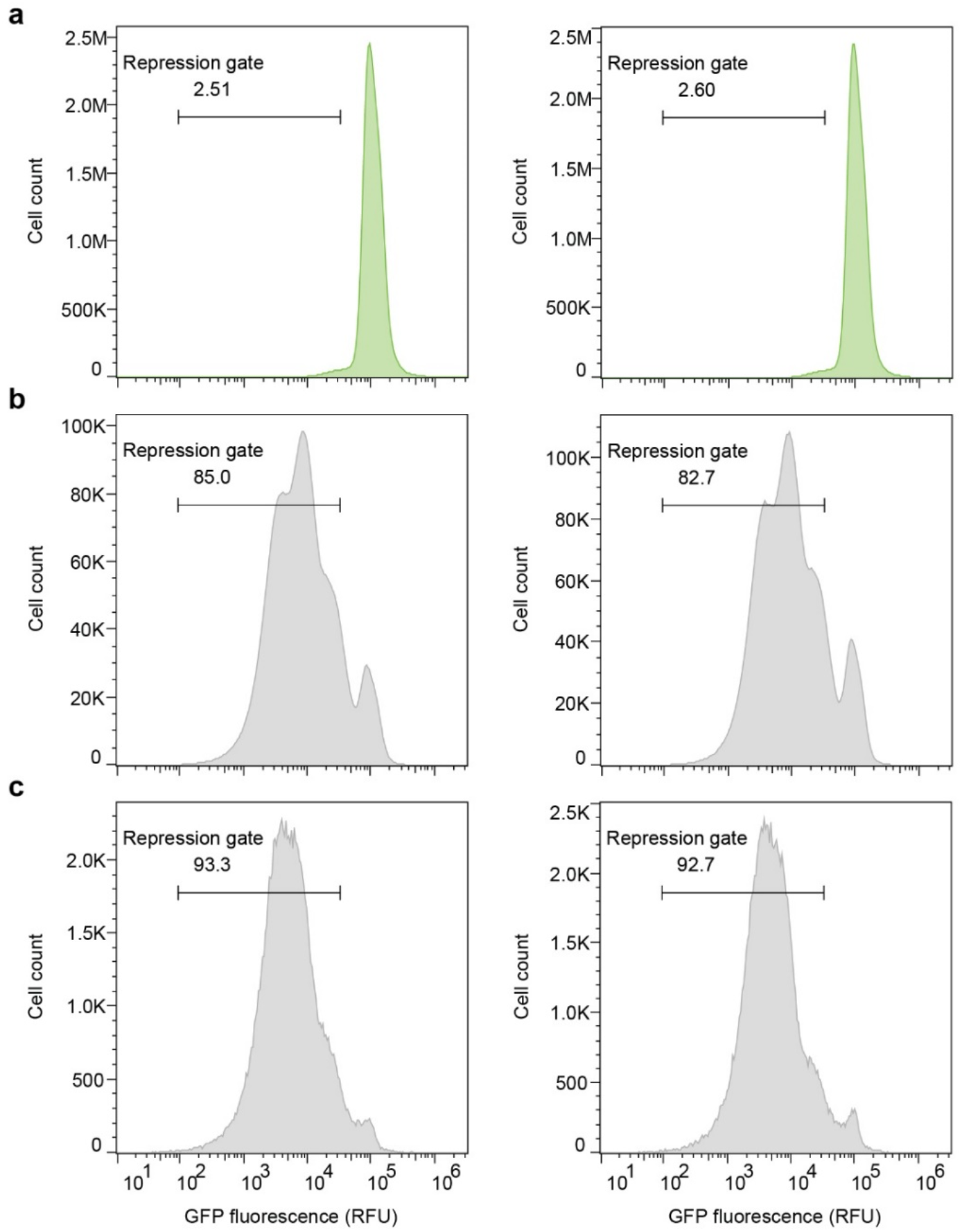
Isolation of repression competent extant and ancestral DBDs using FACS. (**a**), Distribution of GFP fluorescence of the pre-selected phylogenetic DBD library using flow cytometry. The small fraction (2.5-2.6%) of low-fluorescence cells shows that relatively few extant and ancestral DBDs library members repress LacO. (**b-c**), Distribution of GFP fluorescence after one (**b**) or two (**c**) rounds of FACS-based selection to isolate the low fluorescence subpopulation. The indicated repression gate was used to enrich for repression competent variants. RFU, relative fluorescence units.

**Supplementary Fig. 5.**
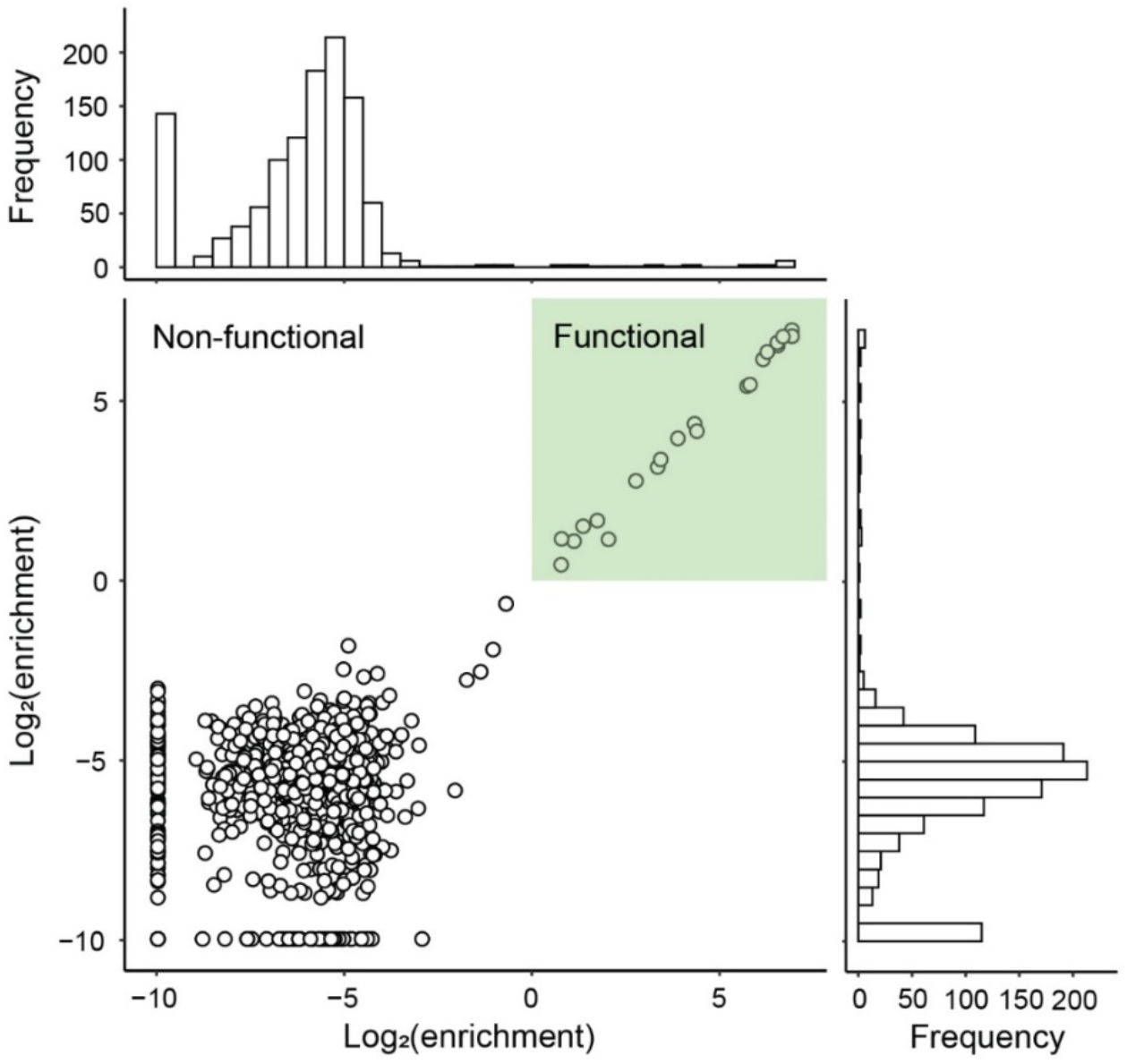
Post-selection enrichment values of extant and ancestral DBD variants. Log transformed enrichment ratios of all 1158 DBD variants as determined by comparing replicate NGS distributions (*n*=2) of the presorted and sorted populations. Higher enrichment values indicate tighter repression of GFP and increased affinity to LacO. Histograms indicate that few (22 of 1158) DBD variants were enriched and functionally repress LacO (green box).

**Supplementary Fig. 6.**
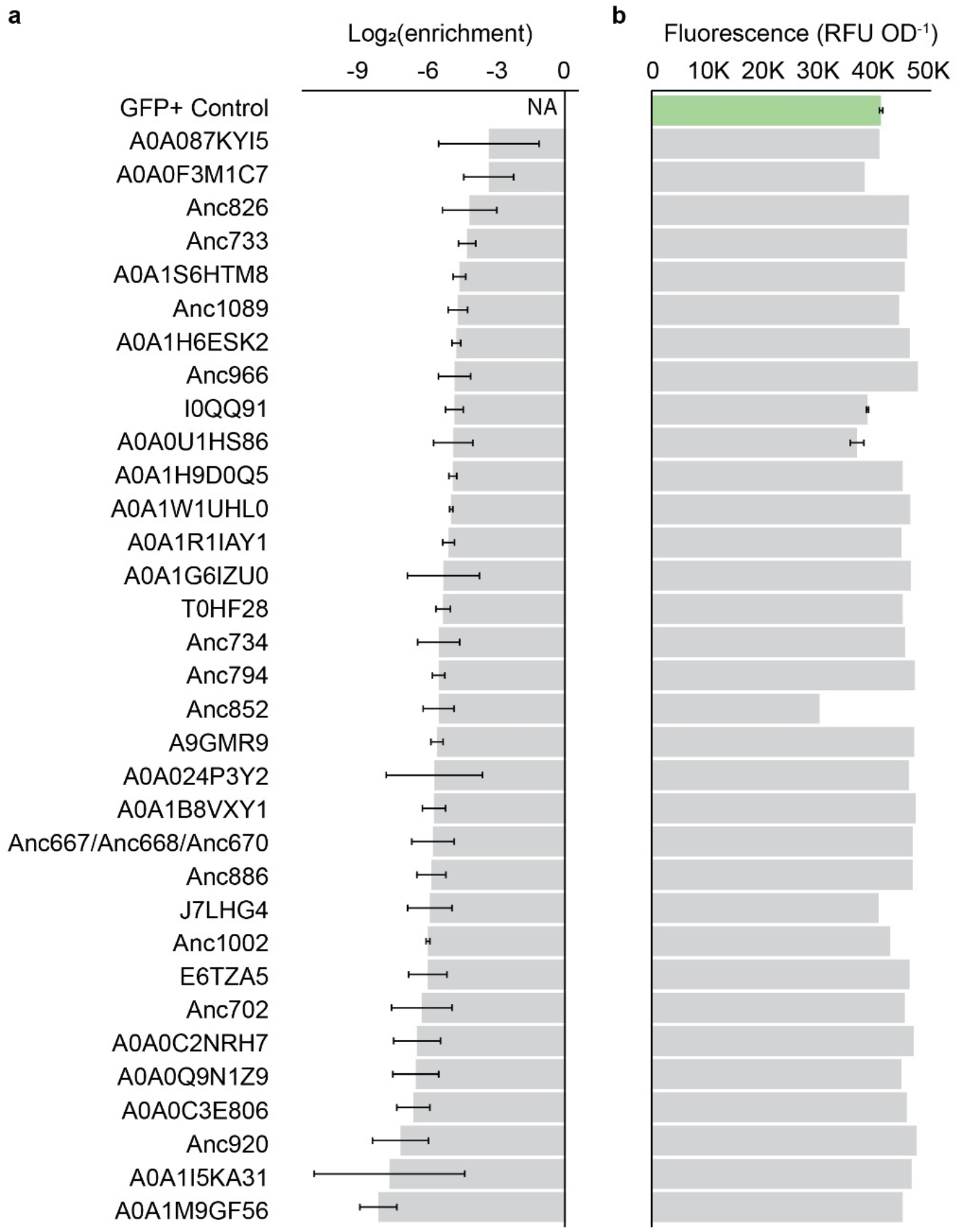
Clonal validation of high-throughput screen. (**a**) Enrichment values of a random subset of depleted ancestral and extant DBD variants derived from the pooled selection and next-generation sequencing experiments. Error bars represent the standard deviation of replicate experiments (*n*=2). (**b**) Clonally assayed mean fluorescence values normalized to OD_600_. Clones were randomly sampled from the pre-selected population and assayed for GFP repression. All depleted clones exhibit high GFP fluorescence (inability to repress LacO) comparable to our GFP+ reporter cells (green). RFU, relative fluorescence units. OD, optical density.

**Supplementary Fig. 7.**
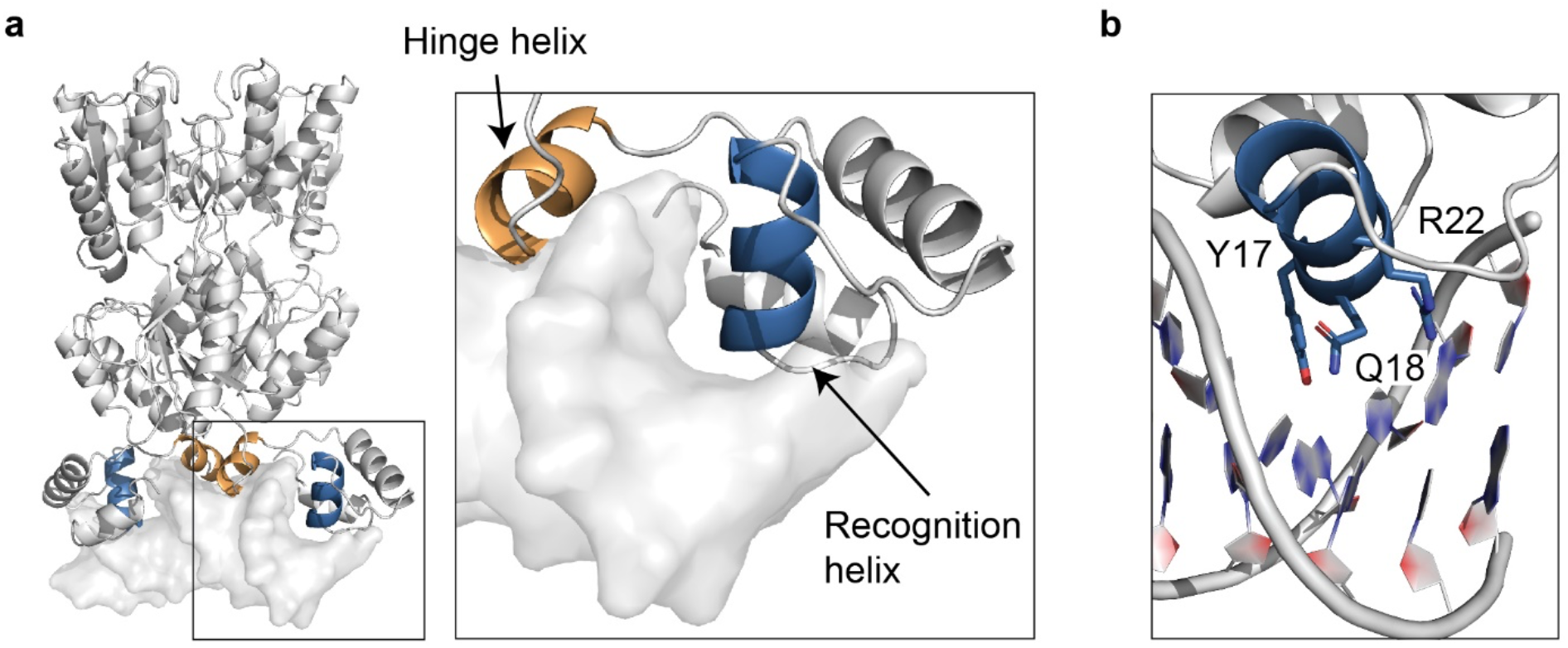
Structural regions of the DBD essential for function. (**a**) Structure (PDB ID: 1EFA) of dimeric extant *E. coli* lac repressor (cartoon) bound to LacO (surface representation). Sequences of functional ancestral and extant DBDs are highly conserved in both the hinge (orange) and recognition helices (blue) (**Fig. 2f**). The hinge helix interacts with the minor groove of LacO, facilitates homodimerization and is involved in allosteric signaling between the LBD and DBD. Functional repressors Anc835, Anc850 and Anc851 are heavily mutated in the hinge helix, but do not relieve GFP repression (**Supplementary Fig. 8**) in the presence of inducer (1mM IPTG), suggesting these mutations may disrupt the allosteric signal transduction pathway. (**b**) Residues of the recognition helix (Y17, Q18, and R22) that facilitate LacO recognition. Y17H and Q18M are the only substitutions tolerated at these critical positions among the functional ancestral and extant DBDs.

**Supplementary Fig. 8.**
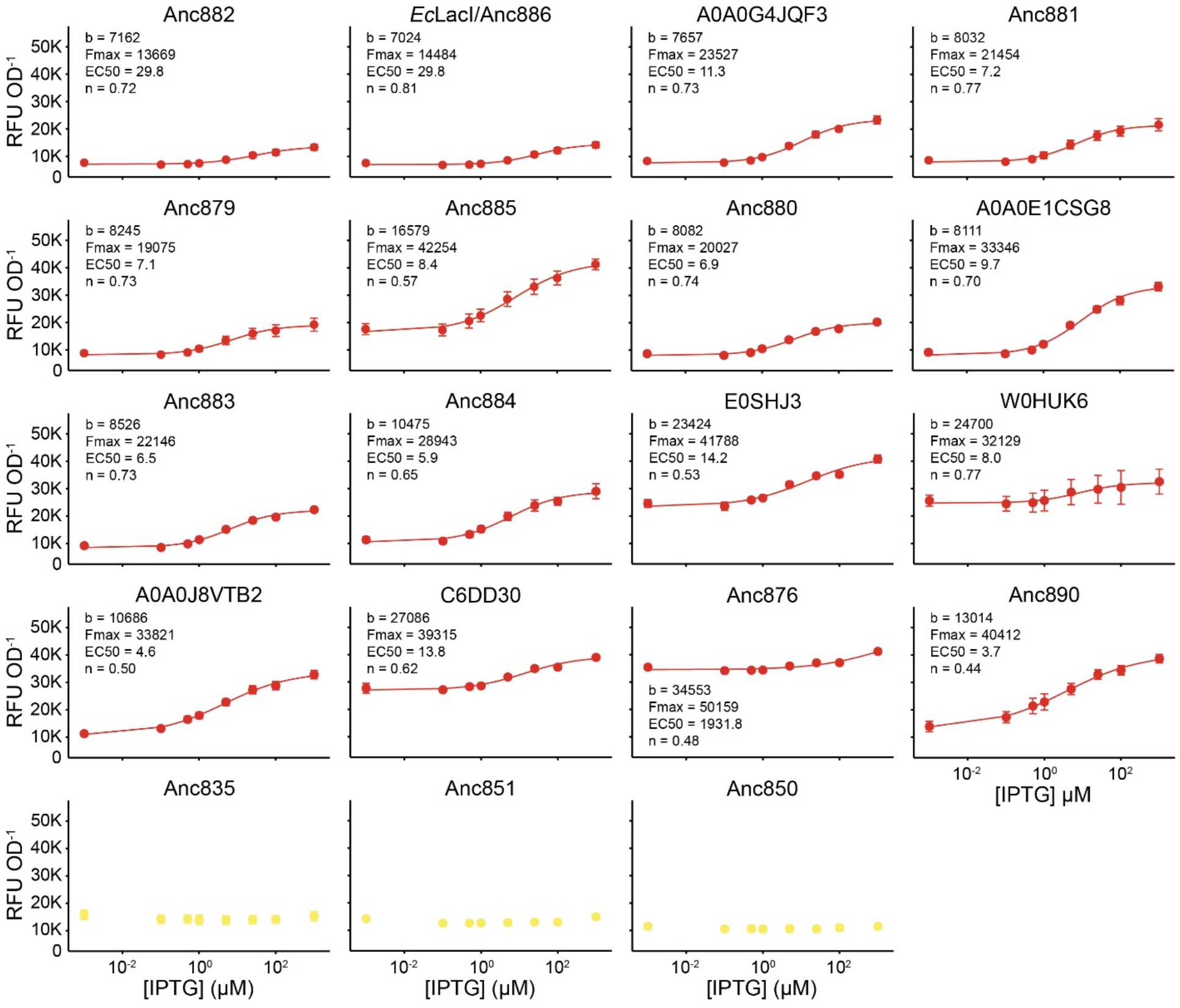
Induction assay of repression competent extant and ancestral DBD variants. The gratuitous inducer IPTG binds to the LBD of *E. coli* LacI to allosterically relieve LacO repression. Dose response curves were fit to the Langmuir-Hill Equation for inducible variants (red). IPTG did not relieve LacO repression for Anc835, Anc851, and Anc850 (yellow). Basal fluorescence, b, is a measure of LacO affinity in the absence of inducer. Fmax is the maximum fluorescence signal achieved at 1mM IPTG. EC50 is the half maximal effective concentration, and n is the Hill coefficient, which is a measure of cooperativity. Error bars represent the standard deviation of replicate experiments (*n*≥2).

**Supplementary Fig. 9.**
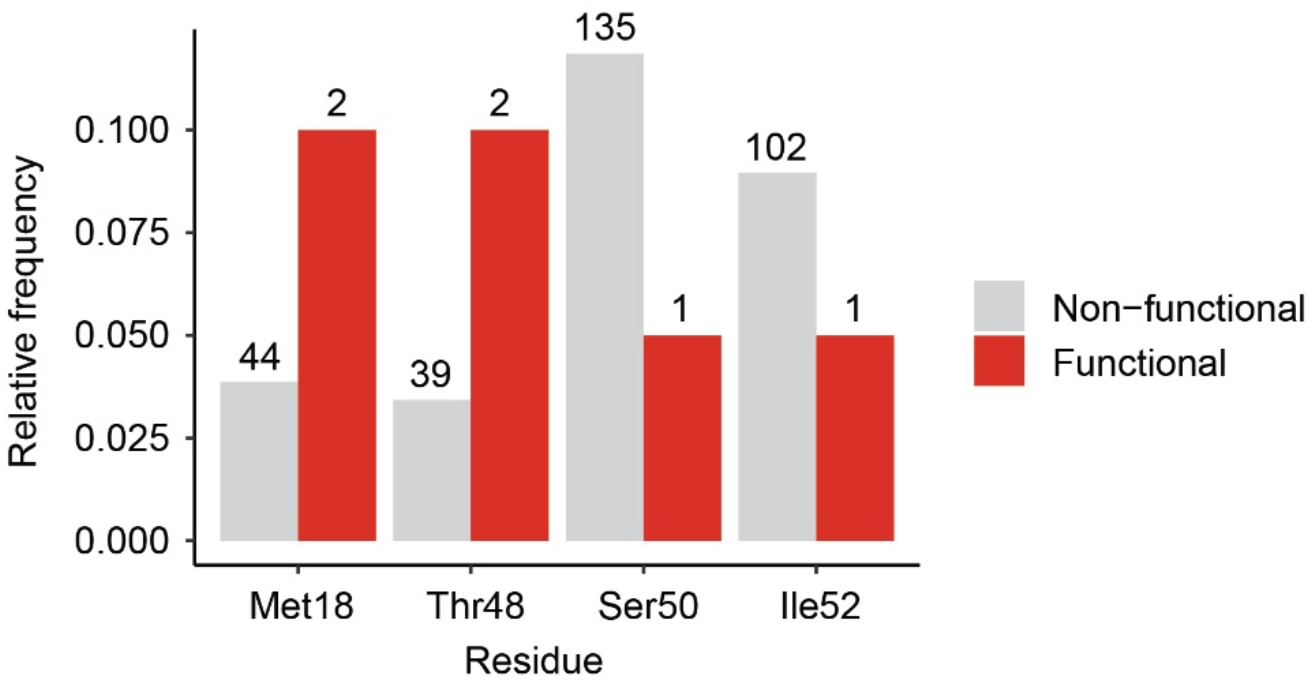
Gln18Met substitution provides additional plasticity for LacO repression. Relative amino acid frequencies of Met18, Thr48, Ser50, and Ile52 for non-functional (gray) and functional (red) phylogenetic DBDs. The Gln18Met substitution in *E. coli* LacI has been previously shown to increase affinity to LacO by 2.5-fold.^39^ Both functional repressors Anc849 and Anc850 have the Gln18Met substitution, and this increased LacO affinity may contribute to the loss of IPTG inducibility (**Supplementary Fig. 8**) and enable less favorable substitutions (Ser50 and Ile52 are more prominent among non-functional DBDs) to be tolerated in Anc851.

**Supplementary Fig. 10.**
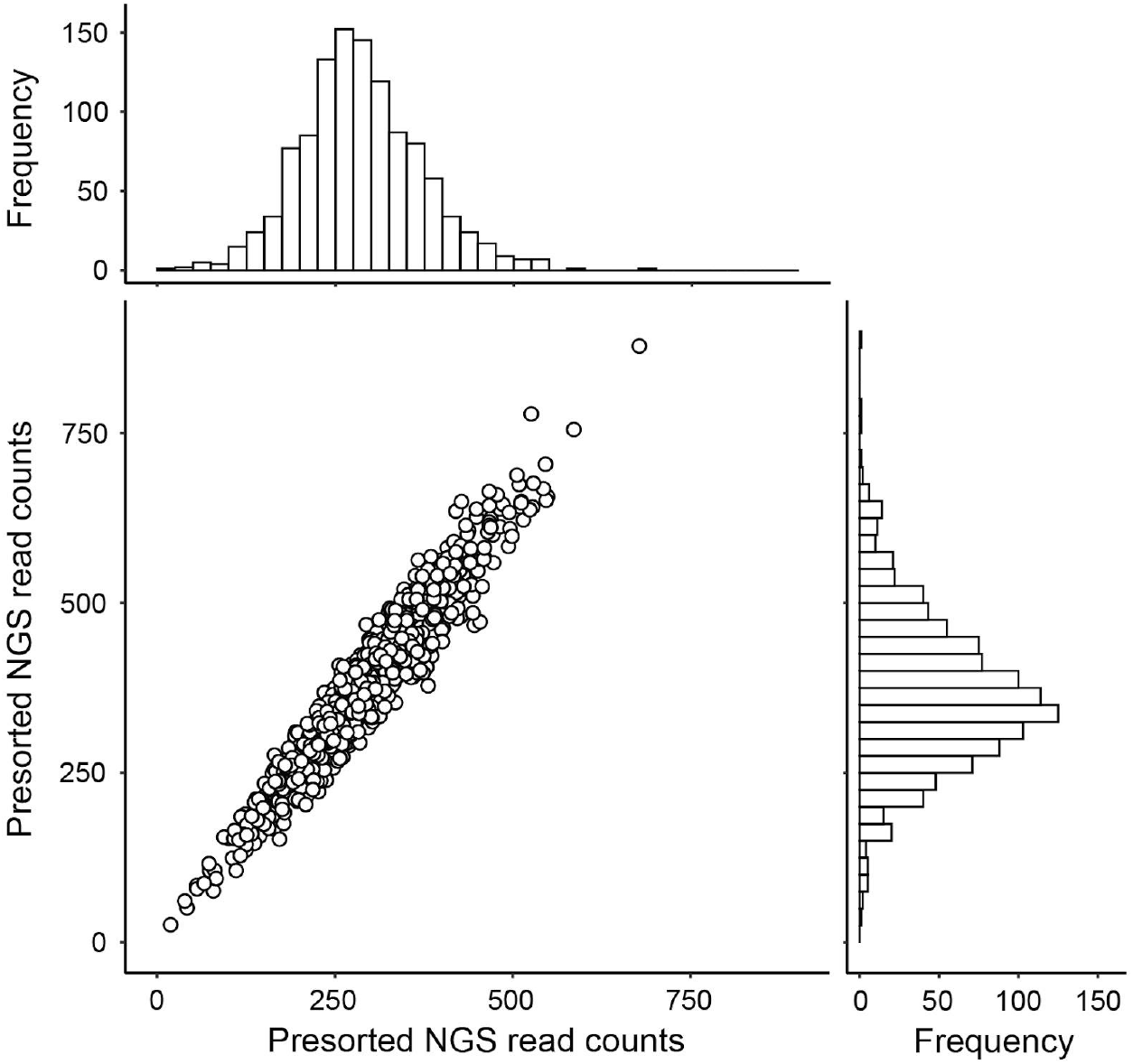
Presorted library distribution of *E. coli* LacI DBD deep mutational scanning variants. Raw read counts of all 1121 DBD single mutant variants from replicate next-generation sequencing runs (*n*=2). All DMS variants were present in the pooled library (median=320±50 reads) prior to selection of functional repressors. Histograms of presorted read counts show that DBD variants were approximately normally distributed with minimal skew (skewness=0.38±0.04).

**Supplementary Fig. 11.**
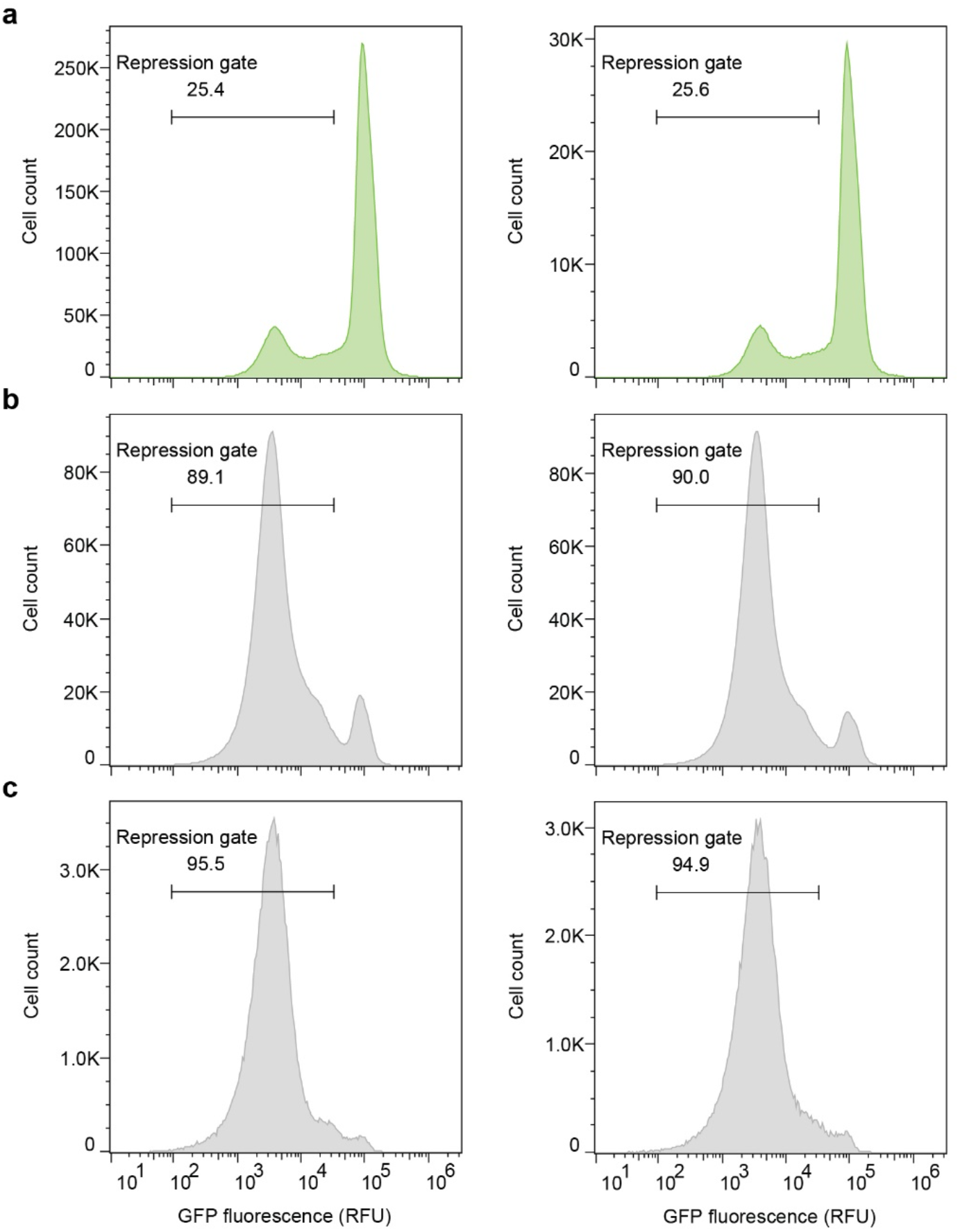
Isolation of repression competent *E. coli* LacI DBD deep mutational scanning variants using FACS. (**a**), Distribution of GFP fluorescence of the pre-selected library using flow cytometry. A larger fraction of low-fluorescence cells (25.4-25.6%) was observed for the DMS library than the phylogenetic library, which indicates that many more DMS variants repress LacO. (**b-c**), Distribution of GFP fluorescence after one (**b**) or two (**c**) rounds of FACS-based selection to isolate the low fluorescence subpopulation. The indicated repression gate was used to enrich for repression competent variants. RFU, relative fluorescence units.

**Supplementary Fig. 12.**
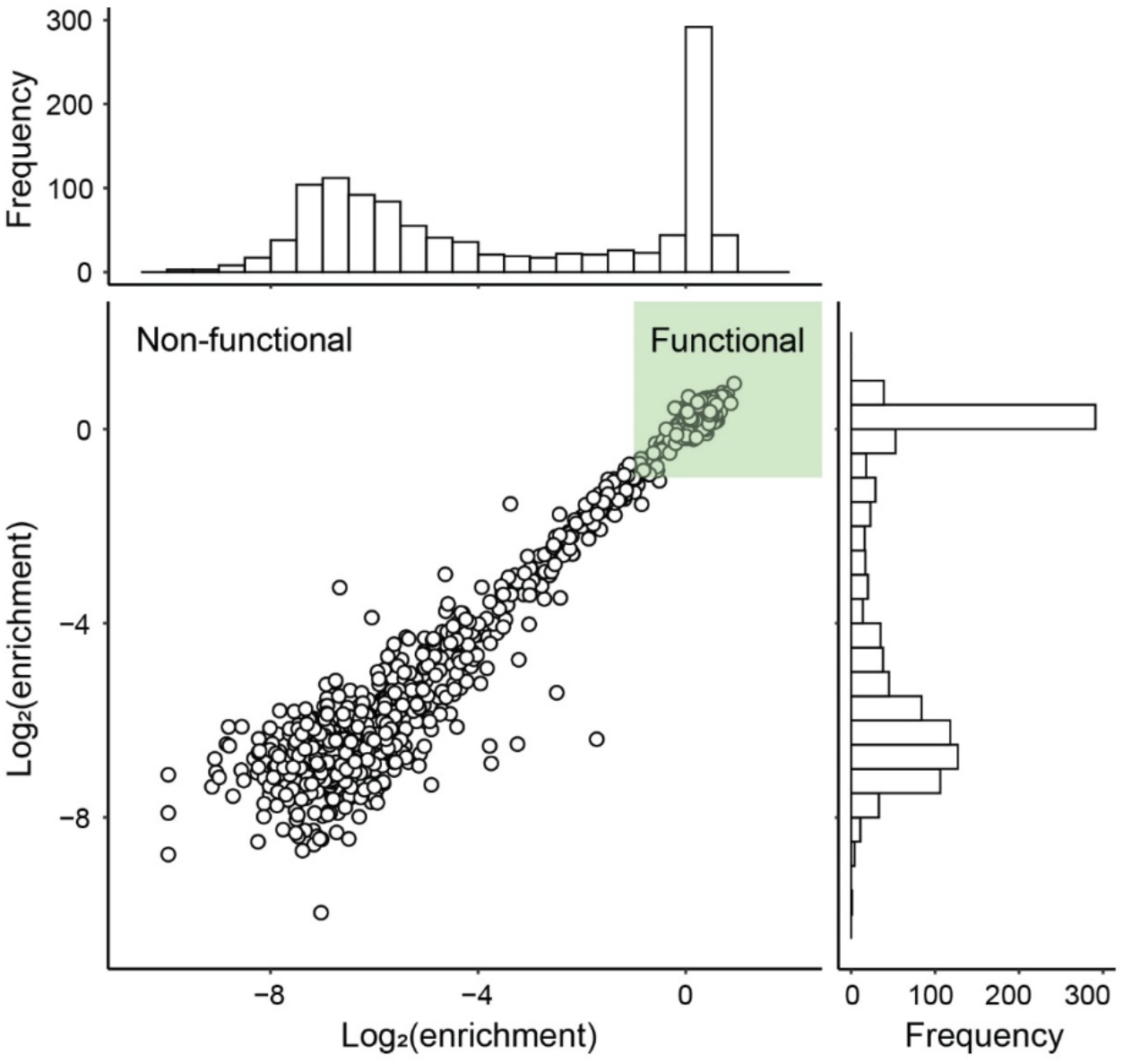
Post-selection enrichment values of *E. coli* LacI DBD deep mutational scanning variants. Log transformed enrichment ratios of all 1158 DBD variants as determined by comparing replicate NGS distributions (*n*=2) of the presorted and sorted populations. Higher enrichment values indicate tighter repression of GFP and increased affinity to LacO. Histograms indicate that 400 (35%) DBD variants were enriched and functionally repress LacO (green box).

**Supplementary Fig. 13.**
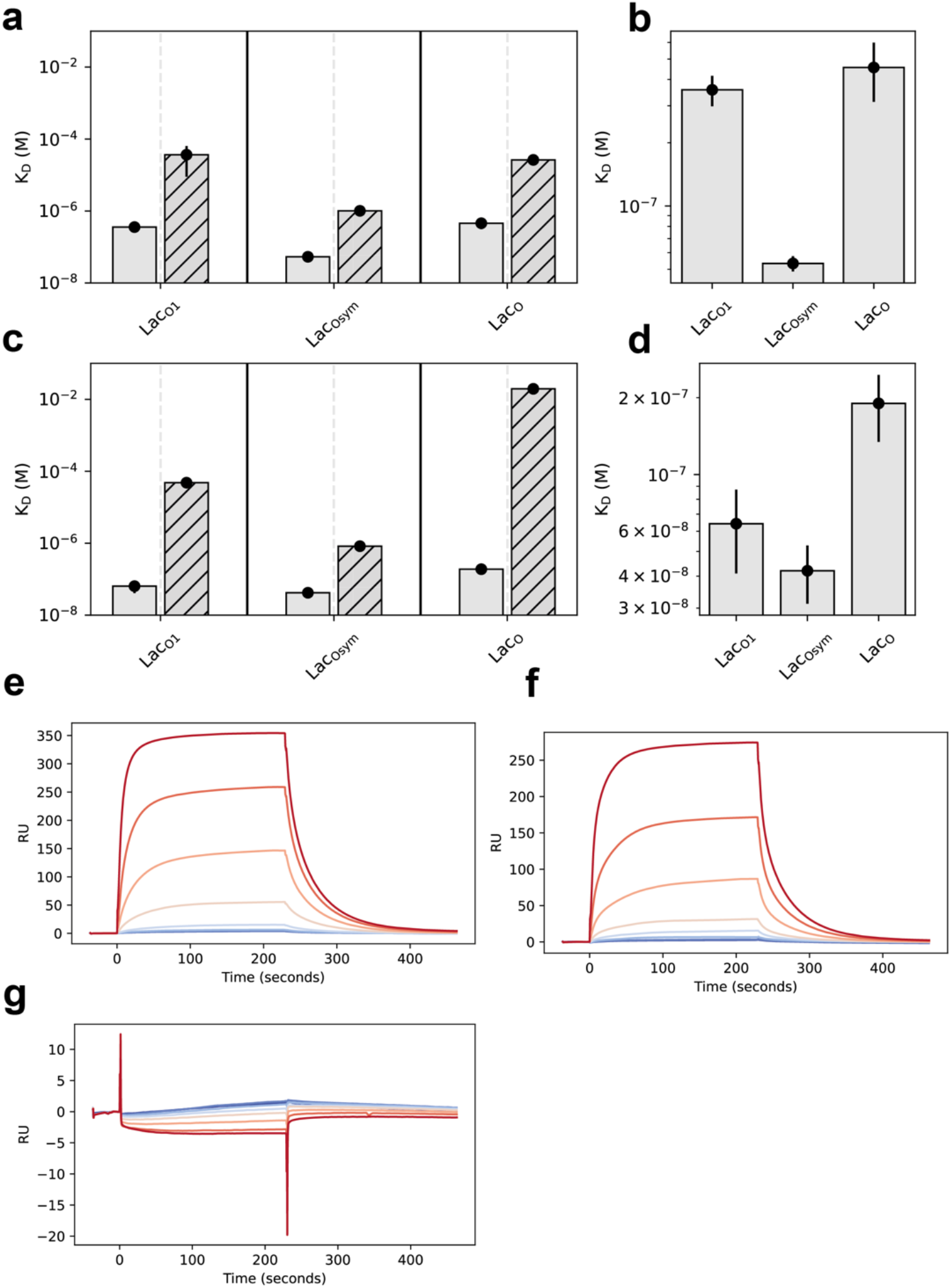
Surface plasmon resonance kinetics and sensograms. (a) K_d_ constants of Anc880 for LacO_1_, LacO_sym_ and LacO when repressed (solid) and induced with 70 μM IPTG (hashed). (b) Zoomed in view of repressed K_d_ constants for Anc880 binding. (c) Kd constants of Anc881 for LacO_1_, LacO_sym_ and LacO when repressed (solid) and induced with 70 μM IPTG (hashed). (d)) Zoomed in view of repressed K_d_ constants for Anc881 binding. (e) Sensogram for Anc880 LacO binding at protein concentrations of 250 nM – 3.91 nM in a log2 dilution series. (f) Sensogram for Anc881 LacO binding at protein concentrations of 250 nM – 3.91 nM in a log2 dilution series. (g) Sensogram for Anc882 binding random DNA at protein concentrations of 250 nM – 3.91 nM in a log2 dilution series. There is no detectable binding for random DNA.

**Supplementary Fig. 14.**
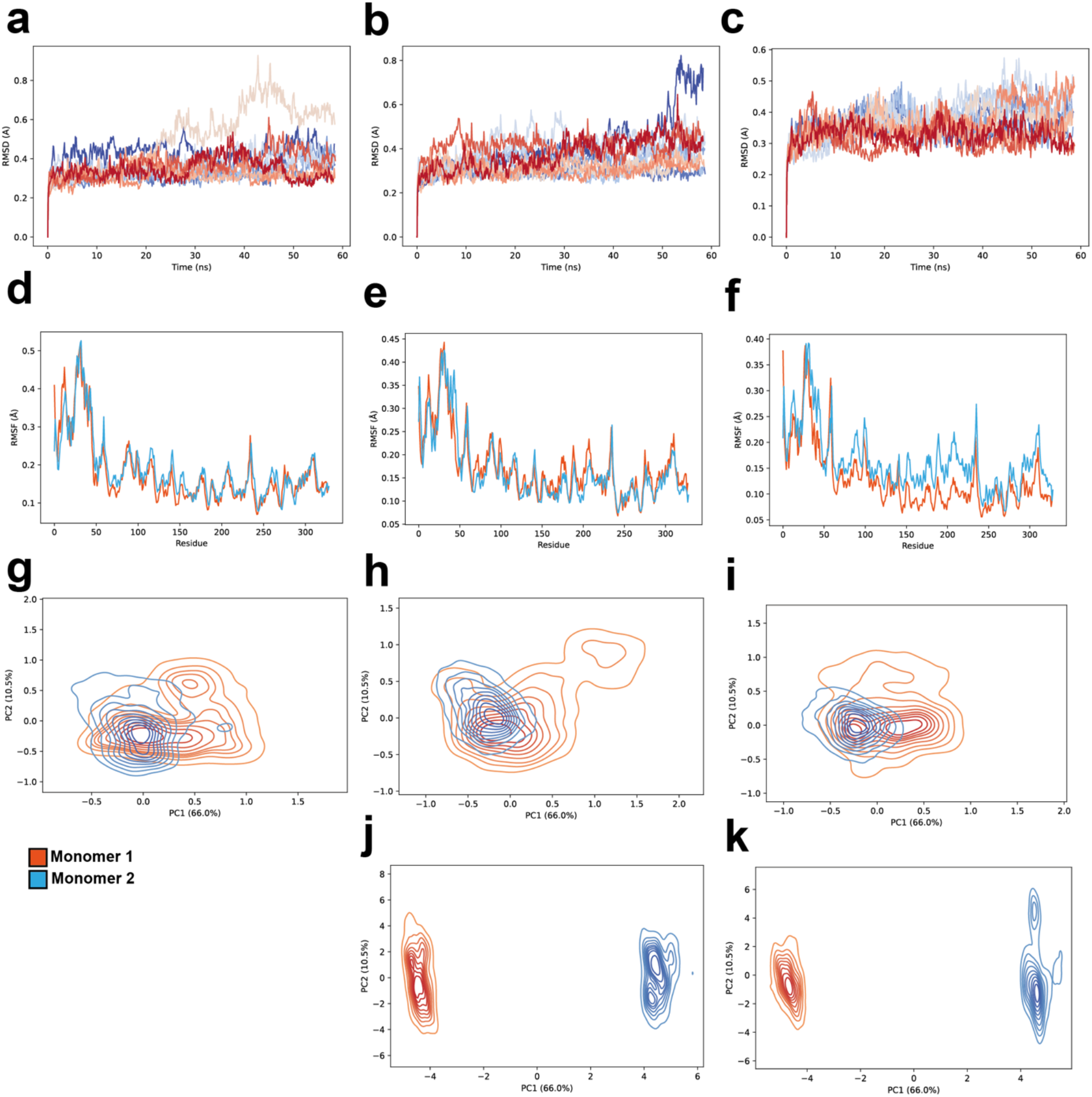
Molecular dynamics simulations of Anc880, Anc882 and EcLacI. (a – c). RMSD evolution of Anc882, Anc880 and EcLacI, respectively. The 10 independent replicates are distinguished by colour. (d-f) RMSF split by monomer for Anc882, Anc880 and EcLacI, respectively. (g-i) PCA of backbone alpha carbons within only the DBD for Anc882, Anc880 and EcLacI, respectively. (j-k) PCA of backbone alpha carbons over the full regulator structure for Anc880 and EcLacI.

**Supplementary Fig. 15.**
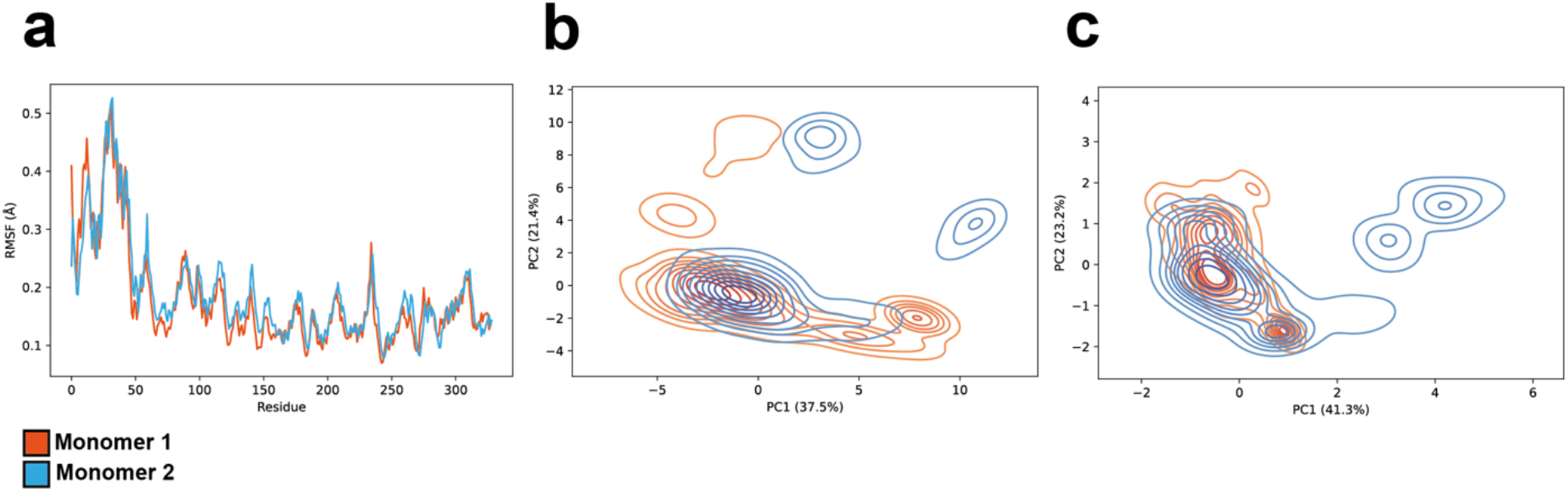
Molecular dynamics simulations of apo-Anc882. (a) RMSF of apo- Anc882 trajectories split by monomer. (b) PCA of backbone alpha carbons over the full apo-Anc882 structure. (c) PCA of backbone alpha carbons over the apo-Anc882 DBD. Motions between monomers in the apo-Anc882 trajectory are highly similar.

